# Molecular and spatial profiling identifies immune endotypes for the stratification of OA patients

**DOI:** 10.64898/2026.01.10.698810

**Authors:** Nicolas Gaigeard, Anaïs Cardon, Romain Guiho, Cankut Cubuk, Astrid Ouattara, Célia Lamothe, Jordan Brouard, Julien De Lima, Manzoor Ahmed, Mathilde Le Mercier, Anaïs Defois, Lucie Danet, Jimmy Perrot, Nazim Benzerdjeb, Claire Vinatier, Richard Danger, Laurence Delbos, Sophie Brouard, Nicolas Degauque, Simona Pagliuca, Myles Lewis, Liliane Fossati-Jimack, Alessandra Nerviani, Denis Waast, Benoit Le Goff, Frédéric Blanchard, David Moulin, Costantino Pitzalis, Jérôme Guicheux, Marie-Astrid Boutet

## Abstract

Osteoarthritis (OA) is a prevalent and heterogeneous joint disease in which synovial inflammation drives structural progression and pain. Despite the recognized heterogeneity of OA, the cellular and molecular organization of synovial tissue remains poorly characterized and defining distinct histological and immune endotypes could guide precision medicine and therapeutic targeting. We show that histologically defined synovial pathotypes are conserved across independent cohorts and correspond to distinct molecular immune endotypes. Integration of bulk and spatial transcriptomics with proteomics revealed niche-specific gene and protein signatures, reflecting the anatomical and functional diversity of OA synovium. The lympho-myeloid pathotype was characterized by mature ectopic lymphoid structures containing CD21^+^CD23^+^ follicular dendritic cells, spatially organized T and B cell zones, and clonally expanded T and B cells with shared immune cell receptor motifs, consistent with local adaptive immune activity correlating with radiological joint damage. These findings highlight how immune organization and cellular composition shape OA pathogenesis and provide a framework for endotype-guided stratification and therapeutic targeting.

## INTRODUCTION

Osteoarthritis (OA) affects approximately 600 million people worldwide, and the OA cases frequency has increased of over 130% since 1990, imposing a substantial socio-economic burden^1^. Historically described as a degenerative “wear and tear” condition associated with aging, OA is now recognized as a complex and heterogeneous inflammatory disease, involving multiple pathophysiological mechanisms beyond cartilage degradation, osteophyte formation, and subchondral bone remodeling. In OA, a persistent low-grade synovial inflammation, fueled by innate and adaptive immune responses, plays a central role in driving joint damage^2^. Although promising strategies have emerged and preclinical studies have shown encouraging results, there is still no effective disease-modifying treatment for OA^3^. The inability to identify patient subgroups more likely to respond to chondroprotective or anti-inflammatory drugs may explain the repeated failure of clinical trials in OA^4^.

The concept of OA phenotypes, defined by clinical, imaging, or etiological features, has emerged to better reflect disease heterogeneity. Recognized phenotypes include patients with predominant pain, metabolic dysregulation, or inflammatory features^5,6^. As also observed in other joint diseases, such as rheumatoid arthritis (RA) or psoriatic arthritis (PsA), the identification of clinical OA phenotypes has improved our understanding of disease heterogeneity, but these phenotypic categories remain insufficient to guide precision medicine^7^. Such observation highlights the need to move beyond descriptive phenotypes toward the definition of disease endotypes, subgroups of patients characterized by specific molecular, cellular, and immunopathological pathways. The definition of disease endotypes has the potential to bridge clinical presentation with pathogenic mechanisms, enabling the identification of predictive biomarkers and the development of targeted, mechanism-based interventions. In OA, where heterogeneity in affected tissues composition is now recognized, the shift from phenotype-based to endotype-driven classification represents a critical step toward relevant personalized therapeutic strategies^5^. While multiomic data from easily accessible biofluids such as plasma, synovial fluid or urine have proven useful for patient classification^8–10^, incorporating changes within joint tissues, readily obtainable from rheumatic patients^11^, may provide additional value for stratification.

Among the joint tissues affected by OA, synovium has gained increasing attention as a central driver of disease pathogenesis. Synovial inflammation (synovitis) is detected in a substantial proportion of patients, correlates with pain, cartilage damage, and structural progression, and can precede radiographic signs of disease^2,12^. OA synovium exhibits variable degrees of immune cell infiltration, involving both innate and adaptive cells^13^. While macrophages are prominent and play a central role^14^, growing evidence highlights the contribution of lymphocytes. As we previously demonstrated^13^, these cells can form organized aggregates called ectopic lymphoid structures (ELS), in over half of OA patients’ synovial tissues at late stage of the disease. Notably, the infiltration and polarization of T cells occur within the OA synovium from early stages of OA^15^, and CD4 deficiency has been shown to reduce disease development in a post-traumatic murine model of OA^16^. However, T cell receptor (TCR) repertoire has largely been unexplored in OA^17^. The presence of B cells in OA synovium also suggests an immunological memory response, likely sustained by T cell help, and a potential antigen-specific involvement in perpetuating inflammation and tissue damage^18^. Changes in peripheral blood immune cell composition, and notably in memory T and B cells, have also been observed in OA patients^19^. Interestingly, circulating autoantibodies have been detected in OA patients^20–22^, which strengthen the hypothesis of a potential contribution of the self-reactivity mechanisms to the disease pathogenesis. However, whether B cell activation is a consequence of chronic low-grade inflammation observed in OA or relates to an active and targetable pathophysiological process remains to be clarified.

Recently, we proposed a novel pathological classification of OA based on synovial histological pathotypes^13^, inspired by concepts described in RA, a prototypical synovitis-driven autoimmune disease where pathotypes correlate with clinical phenotypes, disease progression, and treatment response^23–27^. It is now important to validate whether these synovial pathotypes, namely pauci-immune (PI), diffuse-myeloid (DM) and lympho-myeloid (LM), are consistently observed in independent OA cohorts and whether their presence is associated with damage in adjacent tissues (e.g., cartilage, infrapatellar fat pad, subchondral bone). Establishing such associations would underscore the clinical relevance of this classification. Moreover, determining whether histological pathotypes align with molecular endotypes could pave the way for OA patient stratification and the development of precision therapeutic approaches. Yet, a critical dimension remains largely overlooked: the presence and clinical significance of ELS in OA. Filling this knowledge gap is essential for understanding the immunopathological diversity of the disease and for advancing toward targeted, mechanism-based interventions.

Here, we demonstrate that OA synovial pathotypes are robustly conserved across two independent patient cohorts, reinforcing their potential as a clinically meaningful framework. Using a multiomic strategy that integrates bulk RNA sequencing (RNA-seq), protein and whole-transcriptome GeoMx Digital Spatial Profiling (DSP), we provide an in-depth, spatially resolved characterization of synovial tissue changes in OA. Importantly, we show that OA histological pathotypes correspond to distinct immune endotypes, marked by distinct and unique transcriptional programs, and reveal novel molecular targets that could support endotype-based therapeutic strategies in OA. Finally, through single-cell RNA-seq, spatial transcriptomic using CosMx Spatial Molecular Imaging (SMI), and BCR/TCR-sequencing, we dissect the cellular architecture and clonal organization of the lymphocyte infiltrate, providing unprecedented insight into the nature, origin, and functional relevance of adaptive immune activation in OA.

## RESULTS

### Immune cell-defined synovial pathotypes are conserved across OA cohorts and linked to joint damage severity

As previously demonstrated^13^, synovial tissues from OA patients can be categorized into three histopathological subgroups or pathotypes, namely PI, DM and LM, based on the presence and distribution of macrophages, T, B, and plasma cells. Here, we comparatively analyzed two independent cohorts of patients, 1/ Cohort 1 was recently described^13^, and was composed of 186 end-stage OA patients who underwent knee or hip total joint replacement (London, UK), for which we collected synovial tissues; and 2/ Cohort 2 was composed of 132 end-stage OA patients who underwent knee joint replacement surgery (Nantes, France), and for which we collected matched synovial tissues (n=132), tibial plateau cartilage (n=100), and infrapatellar fat pad (n=61) (**Fig. 1A**). Histological and immunohistochemical staining were performed to determine Krenn synovitis scores and pathotypes (Cohorts 1 and 2)^28^, OARSI scores (Cohort 2)^29^, and fat pad level of vascularization, immune cell infiltrate and fibrosis (Cohort 2).

**Fig. 1.**
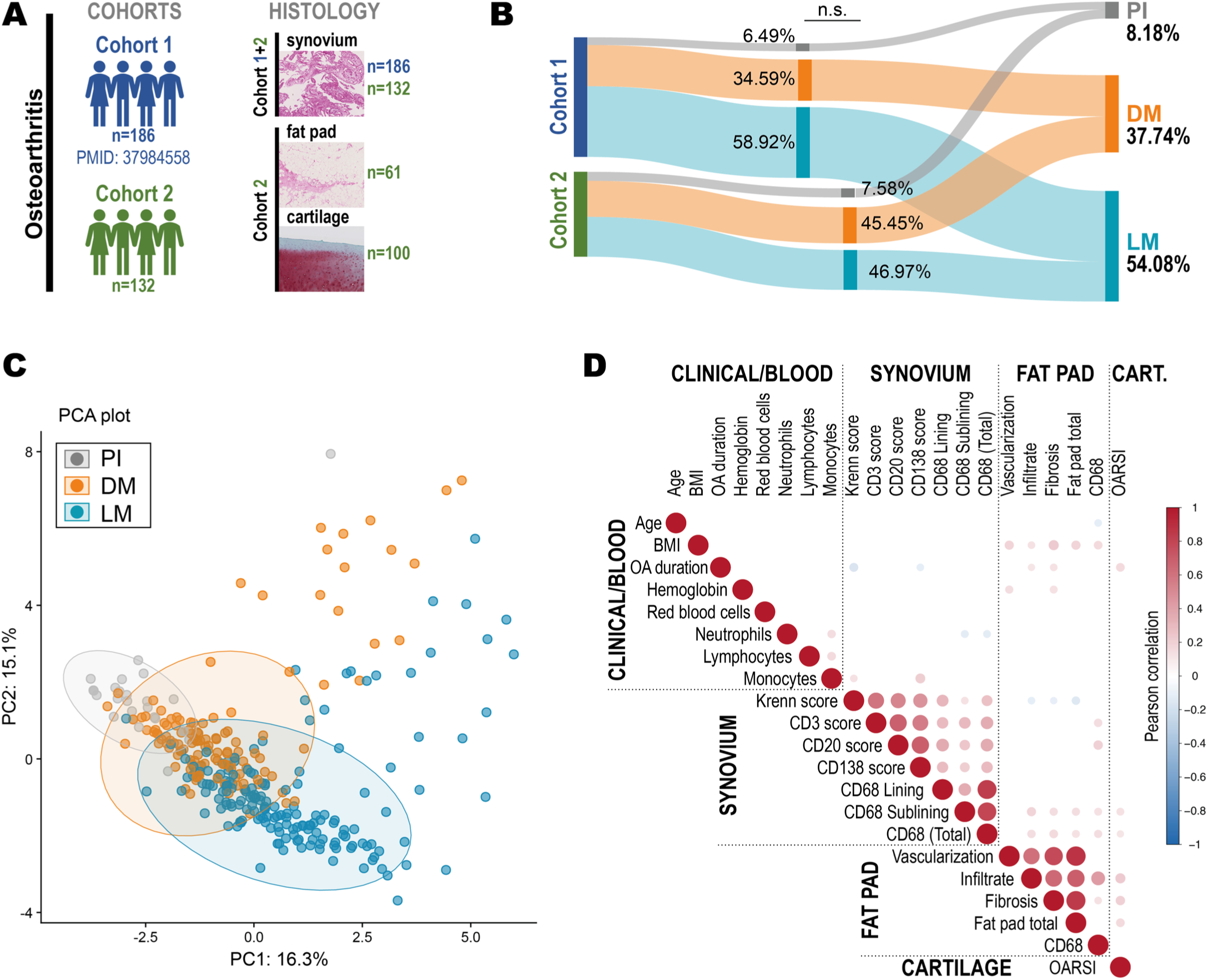
Immune cell-defined synovial pathotypes are conserved across OA cohorts and linked to joint damage severity. **(A)** Cohort 1 was composed of 186 OA patients, and Cohort 2 of 132 patients who underwent knee and hip (Cohort 1) or knee only (Cohort 2) total joint replacement surgery. Synovial tissues (Cohort 1: n=186 and Cohort 2: n=132), tibial plateau (Cohort 2, n=100) and infrapatellar fat pad (Cohort 2, n=61) were processed for histology. **(B)** Riverplot illustrating the distribution of pathotypes in both cohorts (Cohort 1: 12 pauci-immune (PI, in grey), 64 diffuse-myeloid (DM, in orange), 110 lympho-myeloid (LM, in blue); and Cohort 2: 14 PI, 56 DM, 62 LM). Generated with SankeyMatic (https://sankeymatic.com/). **(C)** Principal component analysis (PCA) presenting the distribution of all patients from Cohort 1 and Cohort 2, according to pathotype (PI, DM, LM). The PCA was performed using all available clinical and histological variables, ellipses indicate the 95% confidence interval for each group and axes represent principal components 1 and 2, explaining 16.3% and 15.1% of the total variance, respectively. **(D)** Bubble plot representing the correlation matrix of variables. Circles represent Pearson correlation coefficients between variable pairs, with size proportional to correlation strength and color indicating direction (positive in red or negative in blue). Only statistically significant correlations (p<0.05) are shown. The upper triangle of the matrix is displayed. Missing values were imputed using the median of the corresponding variable prior to analysis. BMI, Body Mass Index; CART, Cartilage; CD, Cluster of Differentiation; OARSI, Osteoarthritis Research Society International score.

We showed that the distribution of synovial pathotypes was not significantly different in Cohort 2, as compared to Cohort 1 (**Fig. 1B**), confirming the reproducibility and robustness of this pathotype classification across independent patient populations (overall distribution: PI, 8.18%; DM, 37.74%; LM, 54.08%). As previously demonstrated for Cohort 1, the percentages of CD3^+^ T cells, CD20^+^ B cells, CD138^+^ plasma cells and CD68^+^ macrophages allowed for an accurate distinction between pathotypes in Cohort 2 (**Fig. S1A**). Although significant differences were observed between pathotypes, and as highlighted before^13^, the pathotype and Krenn classification systems were not redundant, as each pathotype group encompassed patients from multiple Krenn categories (**Fig. S1B**). Importantly, considering data collected as part as Cohort 2, we also showed that the presence of each synovial pathotype was independent of time since diagnosis (OA duration, **Fig. S1C**). Integrating data from patients across both cohorts along with clinical and histopathological parameters, we demonstrated that the three synovial pathotypes formed defined clusters on principal component analysis (PCA) plot, despite partial overlap likely due to inherent disease heterogeneity (**Fig. 1C**). The PCA biplot highlighted key variables driving the variance among samples, such as synovial and fat pad histological features (**Fig. S1D**). Correlation analyses, summarized in the bubble plot (**Fig. 1D**), revealed a structured correlation network in which synovial features cluster and fat pad features form a distinct cluster, whereas clinical variables display selective associations with specific histological features, suggesting potential clinical-pathological interplay. Of note, OA duration positively correlated with cartilage damage (OARSI score), but negatively with Krenn score, and a high BMI correlated with fat pad fibrosis and inflammation. Moreover, synovial and fat pad macrophage infiltrate were positively correlated, emphasizing the potential interplay between synovial and adipose compartments in OA.

### OA synovial pathotypes define distinct immune endotypes with specific transcriptomic programs

To further understand whether synovial histological pathotypes could represent relevant endotypes of the disease, we analyzed OA synovial tissues randomly selected within each pathotype, subject to tissue availability and RNA quality, from both cohorts by bulk RNA-seq (Cohort 1, n=79; Cohort 2, n=15; **Fig. 2A** and **Table. S1**). Following successful technical batch effect adjustment (**Fig. 2B**), PCA illustrated group-specific clustering along PC1 (26% of variance) and PC2 (13% of variance), with partial overlap suggesting both distinct and shared underlying patterns (**Fig. 2C**). Consistent with their distinct histopathological features, the PI, DM and LM pathotypes exhibited specific molecular signatures (**Fig. 2D** and **Fig. 2E**). Gene Ontology (GO) analysis (**Fig. S2A**) and single-sample Gene Set Enrichment Analysis (ssGSEA) (**Fig. 2F**) revealed that the PI pathotype was characterized by the enrichment of pathways related to tissue fibrosis and extracellular matrix remodeling, whereas the DM pathotype showed enrichment in angiogenesis- and metabolism-associated processes. In line with the presence of ELS in the LM pathotype, these tissues displayed upregulation of genes involved in T and B cell activation and adaptive immune responses. Based on bulk transcriptomic analyses, we notably identified candidate genes for each pathotype, that may serve as future therapeutic target or biomarker: Asporin (*ASPN*) and Osteoprotegerin (*TNFRSF11B*) for PI, Cluster of Differentiation 34 (*CD34*) and Amine Oxidase Copper Containing 3 (*AOC3*) for DM, and *CD27* and C-X-C chemokine receptor type 4 (*CXCR4*) for LM, which are involved in joint remodeling (PI), angiogenesis and tissue repair (DM), and leukocyte homing and activation (LM), respectively (**Fig. 2G** and **Fig. S2B**). Cell-specific modules were also analyzed, and the predicted cell composition for each pathotype was consistent with its histological features (**Fig. S3A**). This analysis further revealed a positive correlation between fat pad changes and the fibroblastic-like profile of synovial tissue and confirmed a positive correlation between high histological immune cell scores and increased myeloid and lymphocyte populations (**Fig. S3B**).

**Fig. 2.**
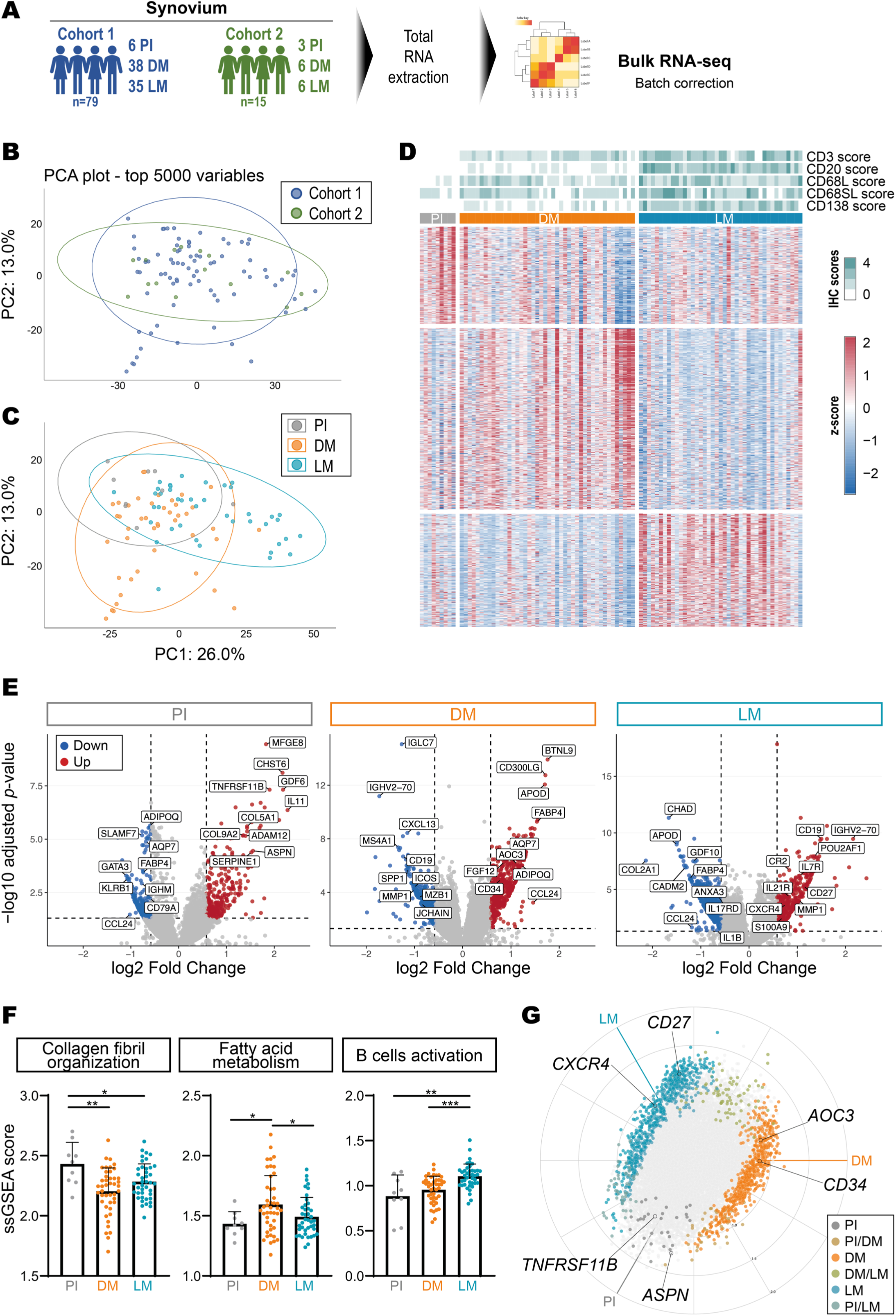
OA synovial pathotypes define distinct immune endotypes with specific transcriptomic programs. **(A)** For bulk RNA sequencing (RNA-seq) analyses, 79 OA synovial samples were analyzed from Cohort 1 (6 pauci-immune or PI, 38 diffuse-myeloid or DM, 35 lympho-myeloid or LM), and 15 from Cohort 2 (3 PI, 6 DM and 6 LM). Total RNA was extracted, bulk RNA-seq was performed and data obtained from both cohorts were considered together following batch adjustment (CombatSeq). **(B, C)** Principal component analysis (PCA) presenting the distribution of selected patients from Cohort 1 and Cohort 2, according to cohort of origin (Cohort 1 in blue, Cohort 2 in green) **(B)** or pathotypes (PI in grey, DM in orange, and LM in blue) **(C)**. Ellipses indicate the 95% confidence interval for each group and axes represent principal components 1 and 2, explaining 26% and 13% of the total variance, respectively. **(D)** Heatmap of differentially expressed genes across the three synovial pathotypes (PI, DM and LM). Each row represents a gene, and each column corresponds to an individual patient. The blue/red scale represents the normalized Z-scores of gene expression. Pathotypes and immunohistochemistry (IHC) scores for T cells (CD3), B cells (CD20), lining macrophages (CD68L), sublining macrophages (CD68SL) and plasma cells (CD138) are annotated above the heatmap. **(E)** Volcano plots presenting differentially expressed genes for each pathotype (PI, DM and LM) assessed by bulk RNA-seq. Each dot represents a gene, the x-axis indicates the –log10(adjusted p-value) and the y-axis indicates the log2(fold-change) in expression between pathotypes. Genes significantly enriched or downregulated in each pathotype, as compared with the two other pathotypes, are highlighted in red and blue, respectively. **(F)** Histograms presenting the single-sample Gene Set Enrichment Analysis (ssGSEA) scores of GOBP_COLLAGEN_FIBRIL_ORGANIZATION, HALLMARK_FATTY_ACID_METABOLISM, and GOBP_B_CELL_ACTIVATION pathways for each pathotype (PI in grey, DM in orange and LM in blue). p-values were calculated using the Kruskal-Wallis test with Dunn’s post-test, * p<0.05; ** p<0.01; *** p<0.001. **(G)** Radial plot prepared from a 3D volcano plot showing differentially expressed genes (in color: p-adjust>0.05) across the three pathotypes (PI, DM and LM). Each dot represents a gene, the distance from the center indicates the log2(fold-change), while the angular position reflects pathotype-specific expression bias. Selected genes of interest are shown. *ASPN*, Asporin; *TNFRSF11B*, TNF Receptor Superfamily Member 11b or osteoprotegerin; *CD34* and *CD27*, Cluster of Differentiation 34 and 27; *AOC3*, Amine Oxidase Copper Containing 3; *CXCR4*, C-X-C chemokine receptor type 4.

### Spatial transcriptomics reveals immune niche-specific programs in OA synovium

As reported in RA, a prototypical chronic and autoimmune joint disease, positional identity plays a critical role in synovial tissue organization^30–32^. To further explore disease- and niche-specific molecular profiles in OA synovium comparatively to RA, we applied 35-plex protein and whole transcriptome atlas (WTA) GeoMx DSP analyses to profile the lining, sublining and ELS, as previously described^33,34^ (**Fig. 3A**). We first confirmed that lining (n=55 ROIs), sublining (n=53 ROIs) and ELS (n=29 ROIs) exhibit distinct protein (**Fig. S4A**) and RNA (**Fig. S4B**) profiles and transcriptomic GO signatures (**Fig. S4C**), consistent with their respective organizational features and functional roles within the synovium. Interestingly, the overall transcriptomic signatures of OA and RA synovial tissues overlapped (**Fig. S4D**).

**Fig. 3.**
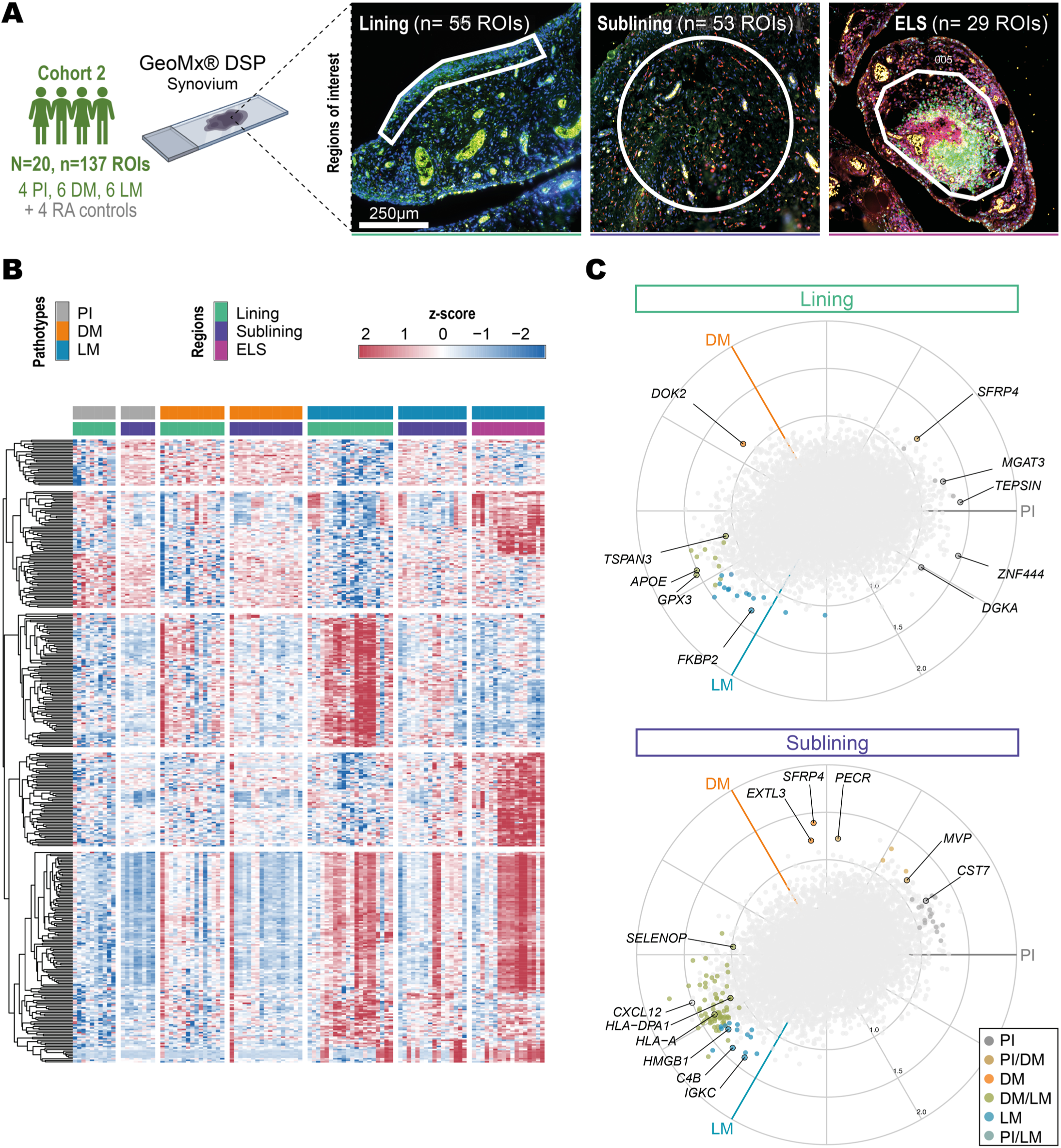
Spatial transcriptomics reveals immune niche-specific programs in OA synovium. **(A)** Representative images of the GeoMx digital spatial profiler (DSP) experiment, including the selection of three regions of interest (ROIs): lining (characterized by the presence of CD68^+^ cells); sublining (characterized by the absence of CD3^+^ and CD20^+^ cells aggregates); ectopic lymphoid structures or ELS (characterized by the presence of CD3^+^ and CD20^+^ cells forming aggregates). Scale bar: 250µm. DSP was performed on 16 OA synovial tissues, belonging to the three pathotype groups: 4 pauci-immune (PI) (10 lining, 8 sublining ROIs), 6 diffuse myeloid (DM) (15 lining, 17 sublining ROIs), and 6 lympho-myeloid (LM) (20 lining, 16 sublining, and 17 ELS ROIs). Synovial tissues from 4 patients with rheumatoid arthritis (RA) were also included (10 lining, 12 sublining, and 12 ELS ROIs). A 35-plex protein and a whole transcriptome panel were applied. **(B)** Heatmap of differentially expressed genes across the three synovial pathotypes (PI, DM and LM) and three synovial niches (lining, sublining and ELS). Each row represents a gene, and each column corresponds to an individual ROI. The blue/red scale represents the normalized Z-scores of gene expression. Pathotypes and regions are annotated above the heatmap. **(C)** Radial plots prepared from 3D volcano plots showing differentially expressed genes across the three pathotypes (PI, DM and LM) in the lining region (upper panel) and the sublining region (lower panel). Each dot represents a gene, the distance from the center indicates the log2(fold-change), while the angular position reflects pathotype-specific expression bias. Selected genes of interest are shown. APOE, Apolipoprotein E; C4B, Complement C4B; CST7, Cystatin F; CXCL12, C-X-C chemokine ligand 12; DGKA, Diacylglycerol kinase alpha; DOK2, Docking protein 2; EXTL3, Exostosin-like glycosyltransferase 3; FKBP2, FK506 binding protein 2; GPX3, Glutathione peroxidase 3; HLA-A, Major histocompatibility complex class I, A; HLA-DPA1, Major histocompatibility complex class II, DP alpha 1; HMGB1, High mobility group box 1; IGKC, Immunoglobulin kappa constant; MGAT3, Mannosyl (beta-1,4-)-glycoprotein beta-1,4-N-acetylglucosaminyltransferase; MVP, Major vault protein; PECR, Peroxisomal trans-2-enoyl-CoA reductase; SELENOP, Selenoprotein P; SFRP4, Secreted frizzled-related protein 4; TSPAN3, Tetraspanin 3; TEPSIN, Clathrin adaptor protein TEPSIN; ZNF444, Zinc finger protein 444.

Spatial RNA profiling confirmed the presence of pathotype-specific signatures initially identified by bulk RNA-seq. Importantly, it also revealed that the molecular profiles of the lining and sublining niches were distinct for each pathotype, and with the ELS niche exhibiting a unique profile specific to the LM pathotype (**Fig. 3B** and **Fig. S5A-B**).

While the PI lining displayed a transcriptomic signature associated with the activation of innate immune receptors typically linked to antiviral and antifungal responses (such as Dectin- or TLR2-dependent pathways, recognizing β-glucans and mannans; including *DGKA*), the LM lining was enriched for pathway signatures associated with cellular stress responses, including *FKBP2*. Genes overexpressed in both DM and LM lining, such as *APOE* and *TSPAN3*, are involved in immune modulation. In contrast, *DOK2*, a negative regulator of tyrosine kinase receptor signaling in lining fibroblasts, was specifically overexpressed in DM lining cells, potentially modulating fibroblast-driven inflammatory and tissue remodeling responses. In the sublining, the PI pathotype was marked by enrichment for innate immune response genesets (including *CST7*, *MVP*), the DM pathotype displayed features associated with stromal cell differentiation, proliferation, and stress responses (including *EXTL3*, *SFRP4*, *PECR*), whereas the both the DM and LM pathotypes were notably enriched for pathway signatures related to natural killer (NK) cell cytotoxicity and T cell-derived mediators production (such as *CXCL12*, *SELENOP*). More specifically, the LM pathotype sublining was characterized by genes associated with the presence of active B cells, plasma cells, and immune stress signals, including *CKB*, *IGKC*, and *HMGB1* (**Fig. 3C** and **Fig. S5C**). The predicted cell composition for each pathotype was consistent with its transcriptomic and histological features (**Fig. S6A**), and the relevance of gene targets highlighted by bulk RNA-seq for each pathotype was confirmed by whole transcriptome DSP (**Fig. S6B**). In particular, the expression of each identified target was significantly enriched in a tissue compartment specific manner, either within the sublining or the ELS (**Fig. S6C**).

Notably, within ELS regions, variable transcriptional profiles across samples were observed (**Fig. 3B**), suggesting heterogeneity in ELS organization and functional state. This variability prompted a more detailed investigation of ELS maturation in LM OA synovium.

### Mature ELS are present in lympho-myeloid OA synovium

Since OA is not classically defined as an autoimmune disease, definitive proof of adaptive immune involvement is still scarce but increasingly suggested^2^. To further understand whether ELS present in LM synovial tissue may exhibit germinal center (GC)-like organizational and functional features, we first assessed CD21 and CD23 protein expression, reflective of the presence of mature follicular dendritic cells (FDC)^35^, in histological synovium sections from 62 LM OA and 16 LM RA as controls (**Fig. 4A**). CD21^-^CD23^-^ and single positive tissues were classified as immature LM (iLM), while synovial tissues exhibiting both CD21 and CD23 expression in the ELS cells were classified as mature LM (mLM) (**Fig. 4B**). Double-immunofluorescence staining ensured the presence of mature FDC characterized by CD21 and CD23 co-expression (**Fig. 4C**). Thirty four percent of OA LM tissues presented mature CD21^+^CD23^+^ ELS, and the distribution of mature/immature ELS was not significantly different from that observed in RA LM tissues (**Fig. 4D**, p=0.49). Of note, and as in RA (**Fig. S7A**), mLM tissues presented a significantly higher cellular density as compared to iLM tissues (**Fig. 4E**), reflective of their potential immune activity. They also exhibited a higher infiltration of T cells, B cells, plasma cells and sublining macrophages (**Fig. S7B**).

**Fig. 4.**
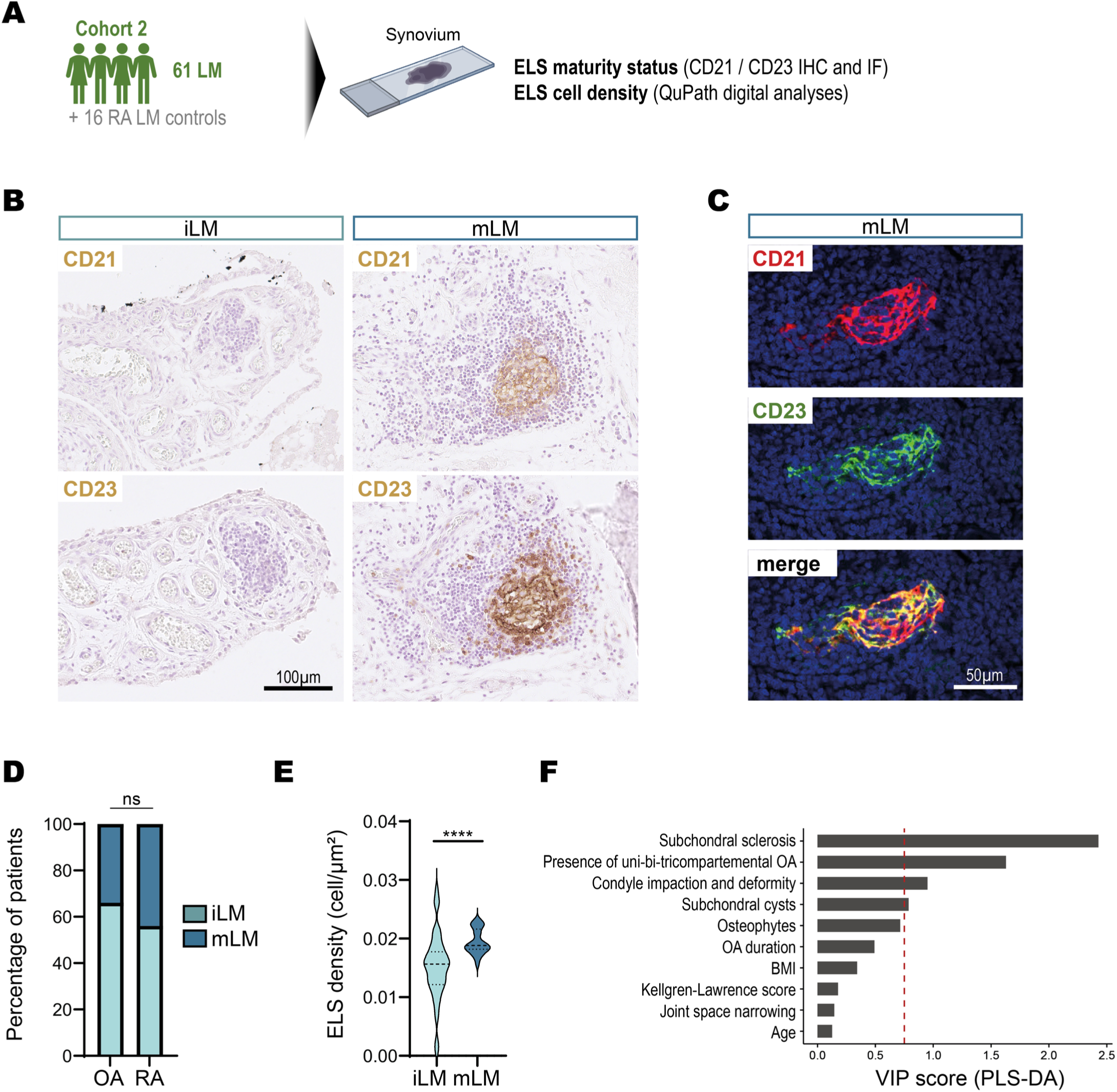
Mature ectopic lymphoid structures (ELS) are present in lympho-myeloid (LM) OA synovium. **(A)** For immunohistochemistry (IHC), immunofluorescence (IF) and ELS cell density status analyses, 61 OA synovial samples were analyzed from Cohort 2 (all lympho-myeloid or LM) and 16 rheumatoid arthritis (RA) synovial tissues were used as controls. **(B)** Representative images of CD21 and CD23 immunohistochemistry (IHC) staining of OA LM synovial tissue sections. Scale bar: 100µm. Immature LM (iLM) are CD21^-^/CD23^-^ or single positive, mature LM (mLM) are CD21^+^/CD23^+^. Representative images of n=40 iLM and 21 mLM samples stained. **(C)** Double immunostaining of CD21 (red) with CD23 (green) in the mLM synovium. Nuclei were counterstained with DAPI (blue). Representative images of n=5 stained samples. Scale bar: 50µm **(D)** Bar chart representing the distribution of iLM and mLM tissues in OA (n= 61) and RA (n= 16). p-value was calculated using the Fisher’s exact test and was not significant. **(E)** Violin plots representing the distribution, median and quartiles values of ELS density in cells/µm^2^ in OA tissues (n= 61 tissues analyzed), assessed using QuPath. Briefly, iLM and mLM ELS areas were manually selected, total cell count was determined using the *Cell Detection* tool and the number of cells was related to the surface of each ELS area. p-value was calculated using the Mann-Whitney test, ****, p< 0.0001. **(F)** Bar plot showing the Variable Importance in Projection (VIP) scores and reflecting the selected features contributing to iLM and mLM group discrimination in the Partial Least Squares Discriminant Analysis (PLS-DA) model. Each bar represents one variable, ranked by decreasing VIP value. The dashed line indicates the threshold of VIP = 0.75, above which variables are considered to contribute meaningfully to the model. BMI, Body Mass Index.

To test whether mature ELS presence marks a clinically severe OA endotype, we analyzed radiographic joint damage in patients with a LM synovial pathotype. Interestingly, Partial Least Squares Discriminant Analysis (PLS-DA) indicated that radiological markers of joint damage (e.g., subchondral sclerosis, cysts, and condylar impaction) mostly contributed to iLM and mLM group discrimination, while other clinical data such as age or time since diagnosis (OA duration) were not involved (**Fig. 4F**). These findings underscore the central role of immune activity, particularly the presence of mature synovial ELS, in shaping the structural severity of OA.

### ELS maturation in lympho-myeloid synovium is associated with a global rewiring of the cellular microenvironment

To comprehensively characterize the cellular and molecular reorganization associated with ELS maturation, we integrated single-cell RNA sequencing and high-plex spatial transcriptomics (CosMx Spatial Molecular Imager, 6,000-gene panel) with advanced spectral flow cytometry analyses (**Fig. 5A**). First, to assess how ELS maturation reshapes the synovial immune ecosystem, we combined the single-cell and CosMx datasets for joint analysis and performed cell clustering and annotation based on literature consensus for stromal, myeloid and lymphoid synovial cells^32,36–38^ (**Fig. 5B** and **Fig. S8A**). UMAP visualization revealed distinct clusters corresponding to major synovial cell populations, including fibroblasts, endothelial and mural cells, macrophages, T and B cells, and plasma cells (**Fig. 5B**). These integrated analyses revealed a marked reorganization of the immune and stromal landscape in mLM synovium, characterized by a reduced proportion of tissue-resident macrophages (TRM) in favor of expanded inflammatory macrophage (IM) subsets, including CLEC10A^+^, S100A12^+^, and SPP1^+^ macrophages. Concomitantly, the proportion of fibroblasts expressing POSTN and CXCL12 was increased, suggesting enhanced stromal activation and chemokine-mediated lymphocyte recruitment. This remodeling was accompanied by an expansion of B lineage cells, including naïve B cells, plasmablasts, and plasma cells, reflecting a reshaping of both innate and adaptive compartments in mature LM synovium (**Fig. 5C**).

**Fig. 5.**
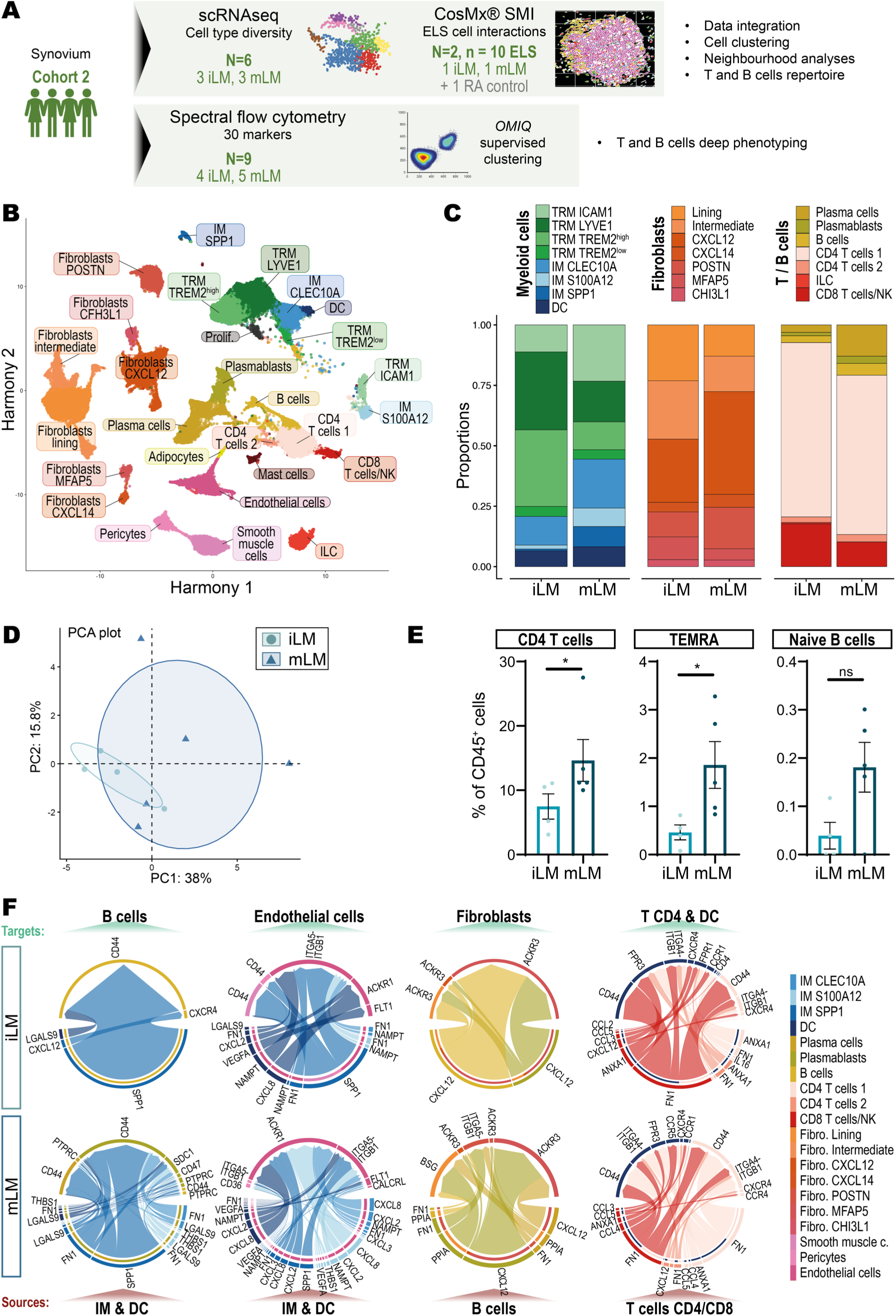
ELS maturation in lympho-myeloid synovium is associated with a global rewiring of the cellular microenvironment. **(A)** Single-cell RNA sequencing (scRNA-seq, N=6, including 3 immature lympho-myeloid iLM and 3 mature LM mLM synovial tissues) and single cell spatial transcriptomic (6000-plex CosMx spatial molecular imager SMI, N=2, including 1 iLM and 1 mLM and 1 rheumatoid arthritis RA tissue as control) data were integrated, as described in the Materials and Methods section and used for cell clustering, neighborhood and TCR/BCR repertoire analyzes. In parallel, spectral flow cytometry was performed on 9 dissociated synovial tissues (4 iLM and 5 mLM) using a 30 markers panel. **(B)** Uniform Manifold Approximation and Projection (UMAP) projection of cells integrated from scRNA-seq and CosMx datasets using Harmony. Each dot represents an individual cell, colored by its cell type identity, as indicated above each group of cells. **(C)** Histogram showing the proportions of each subset of myeloid cells, fibroblasts, or T and B cells in iLM and mLM in the integrated dataset. Each color corresponds to a cell type, as indicated in the legend. CD, cluster of differentiation; CHI3L1, chitinase-3-like protein 1; CLEC10A, C-type lectin domain containing 10A; CXCL, C-X-C motif chemokine ligand; DC, dendritic cells; ICAM1, intercellular adhesion molecule 1; ILC, innate lymphoid cells; LYVE1, lymphatic vessel endothelial hyaluronan receptor 1; MFAP5, microfibril associated protein 5; NK, natural killer cells; POSTN, periostin; S100A12, S100 calcium binding protein A12; SPP1, secreted phosphoprotein 1; TREM2, triggering receptor expressed on myeloid cells 2; TRM, tissue-resident macrophages. **(D)** Principal component analysis (PCA) showing 4 iLM (light blue) and 5 mLM (dark blue) samples. The PCA was performed using the relative abundance (percentage among all CD45^+^ live cells) of each cell subpopulation, defined based on the expression of 30 spectral cytometry markers. PC1 and PC2 explain 38.3% and 15.0% of the total variance, respectively. **(E)** Histograms showing the percentage of CD45^+^ cells represented by CD4^+^ T cells, terminally differentiated effector memory T cells re-expressing CD45RA (TEMRA) cells, and naïve B cells in iLM and mLM samples. p-value was calculated using the Mann-Whitney test, *, p<0.05. **(F)** Chord diagrams generated from CellChat analyses of the integrated dataset illustrating inferred cell-cell communication networks. Upper panels represent iLM samples, lower panels represent mLM samples. The different colors represent the cell identities, as indicated in the legend. Pathways involving inflammatory macrophages (IM), dendritic cells (DC), B cells and CD4/CD8 T cells as sources, B cells, endothelial cells, fibroblasts and CD4 T cells/DC as targets are shown.

Beyond the transcriptomic analyses, we performed deep immunophenotyping of T and B cells by spectral flow cytometry, using the gating strategy described in **Fig. S9A-C**. PCA of 30 markers clearly separated iLM and mLM synovial samples into two distinct groups (**Fig. 5D**), with the iLM group primarily driven by myeloid cell populations and the mLM group mainly dominated by adaptive immune cells (**Fig. S9D**). mLM samples exhibited a selective expansion of CD4^+^ T cells and terminally differentiated effector memory (TEMRA) T cells, mainly CD8^+^, and a trend toward increased naïve B cells, in line with the single-cell observations and providing additional resolution of lymphocyte functional states (**Fig. 5E**). Although flow cytometry did not allow the clear identification of plasma cells, likely due to enzymatic tissue dissociation leading to loss of surface CD38 and CD138 antigens, previous histological analyses demonstrated higher infiltration of CD138^+^ plasma cells, together with B and T cells, in mLM tissues (**Fig. S7B**).

Cell–cell communication inference using CellChat revealed a profound rewiring of intercellular signaling in mLM synovium (**Fig. S8B-C** and **Fig. 5F**). Enhanced interactions were observed between inflammatory macrophages (IM) and dendritic cells (DC) signaling to B and plasma cells (e.g., Thrombospondin-1 [THBS1] to Syndecan-1 [SDC1], involved in matrix remodeling and angiogenesis) and to endothelial cells, pericytes, and smooth muscle cells (e.g., CXCL8/3/2 to Duffy Antigen Receptor for Chemokines [ACKR1], maintaining local chemokine gradients). Interactions were also enhanced between B and plasma cells and lining fibroblasts / inflammatory CXCL12^+^ fibroblasts (e.g., Cyclophilin A [PPIA] to CD147/Basigin [BSG], activating matrix metalloproteinase production), and between CD4^+^ and CD8^+^ T cells with T cells and DCs (e.g., CCL3/4 to CCR5, promoting immune cell recruitment and retention). These interactions are consistent with the formation of a mature, functionally organized immune niche, supporting stromal-immune crosstalk, lymphocyte recruitment and activation.

Together, these data indicate that ELS maturation in LM OA synovium is accompanied by a coordinated remodeling of cellular composition, functional states, and intercellular communication, supporting the emergence of a highly organized and interactive immune microenvironment.

### Mature OA ELS show spatial and cellular features of functional germinal centers

The presence of mature ELS in OA synovium prompted us to further dissect the cellular composition and spatial organization of these structures in iLM, mLM, and in autoantibody-positive RA synovium. Using GeoMx protein and WTA for spatial profiling (**Fig. S10A**), as detailed earlier in this study, we analysed mature and immature OA ELS molecular profiles. Immature ELS displayed signatures linked to innate immune responses, including *IL12B*, whereas mature ELS were enriched for genes associated with plasma cell differentiation, such as *XBP1* and *JCHAIN*, highlighting functional differences between these structures within OA synovium (**Fig. S10B**). Interestingly, at the protein level, KI-67 was notably overexpressed in mature ELS (**Fig. S10C**), indicative of ongoing in situ lymphocyte proliferation. Compared to RA, OA ELS were notable for higher expression of *CCL21* at the transcriptomic level (WTA, **Fig. S10D**), a chemokine produced by lymphatic endothelial and stromal cells that mediates T cell recruitment, whereas no significant differences were observed in protein abundance across the 35-plex panel (**Fig. S10E**).

Using our CosMx SMI dataset, ELS cells were selected using a free-form selection tool (**Fig. S10F**), and the localization of each cell type inside or outside ELS was subsequently confirmed (**Fig. S10G**). Compared with iLM, mLM ELS exhibited organized T and B cell zones (**Fig. 6A**), similar to RA-associated ELS (**Fig. S10H**), suggesting a level of organization consistent with GC-like activity and potential local antigen-driven immune responses. Force-directed graphs representing directed associations between cell types further supported these observations: mLM ELS displayed tight clustering around lymphoid subsets, reminiscent of RA, whereas iLM ELS were organized around macrophages, mural cells, and fibroblasts (**Fig. 6B** and **Fig. S10I**).

**Fig. 6.**
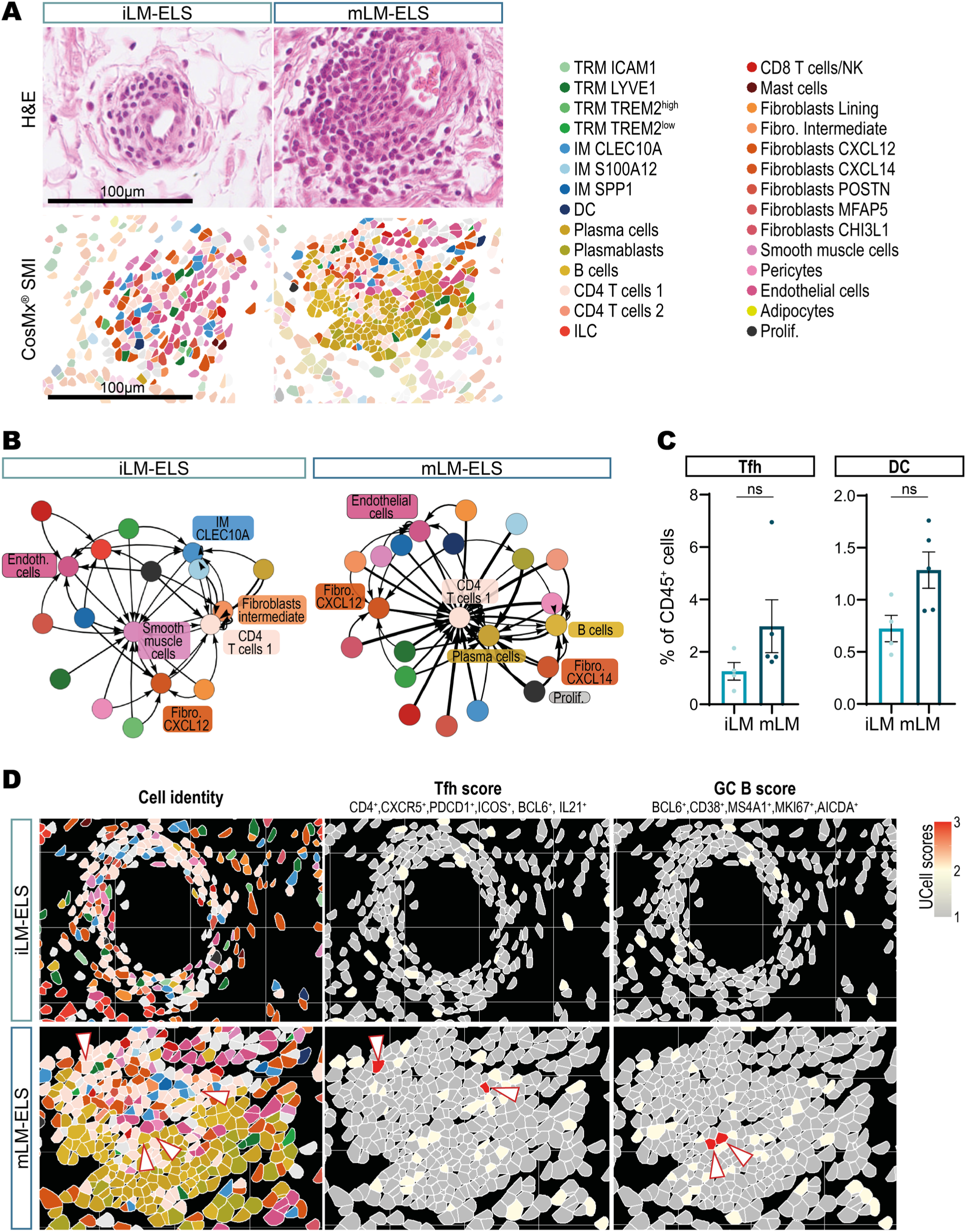
Mature OA ELS show spatial and cellular features of functional germinal centers. **(A)** Representative images of Hematoxilin & Eosin (H&E) and CosMx Image Dimplot representing ectopic lymphoid structures (ELS) cell type composition and organization, each cell type is depicted using a different colour, according to the corresponding legend. Scale bar: 50µm. **(B)** Force-directed graph for immature lympho-myeloid (iLM) and mature lympho-myeloid (mLM) ELS representing directed spatial associations between cell types, derived from the neighborhoods defined by extended Delaunay triangulation (combinatorial radius 4). Each node corresponds to a cell type and is colored accordingly (as specified in the legend in **(A)**). Directed edges (arrows) point from cell type A to cell type B when cells of type B comprise at least 5 % of the neighbors of cells of type A. Edge width reflects the relative weight of these interactions. **(C)** Histograms showing the percentage of CD45^+^ cells represented by follicular helper T cells (Tfh) and dendritic cells (DC) in iLM and mLM samples. **(D)** ImageDimPlot showing cells enriched for UCell AddModule scores of Tfh and germinal center (GC) B cells in iLM and mLM synovium. Cellular identities are colored according to the legend in **(A)**. Gene signatures used to compute each score are indicated. White arrows highlight cells with pronounced enrichment within mature ELS.

Spectral flow cytometry analyses revealed that DC and follicular helper T (Tfh) cell populations were enriched in mLM tissues compared with iLM (**Fig. 6C**). These tissue-level observations were further supported by module-based analyses of the CosMx spatial transcriptomic dataset that revealed the presence of Tfh and GC B cells specifically within mLM ELS, but not iLM (**Fig. 6D**), recapitulating patterns observed in RA (**Fig. S10J**).

Together, these results indicate that mature OA ELS not only exhibit spatial segregation of T and B cells but also display transcriptional and cellular programs characteristic of follicular organization and germinal center activity.

### T and B cell repertoire analysis reveals antigen-driven expansion in mature OA ELS

To further assess the functional maturity of ELS in OA synovium, we analyzed T and B cell receptors (TCR/BCR) immune repertoires comparatively in iLM and mLM samples.

TCR repertoires in mLM samples exhibited a trend toward fewer unique clones, reflecting a potential clonal expansion of certain TCR sequences (**Fig. 7A**). Clonal overlap analysis, using the Jaccard index, showed that TCR repertoires were largely individualized in iLM and mLM samples, with minimal shared clones **(Fig. 7B)**. Interestingly, we observed increased clonal overlap within the mLM subtype, whereas iLM clonotypes showed minimal sharing across patients, suggesting the presence of a shared antigenic stimulus in mLM that may elicit a more organized immune response **(Fig. 7B)**. Importantly, T cell clone expansion in synovium was observed across both CD4^+^ and CD8^+^ populations (**Fig. 7C**). Expanded clones accounted for a substantial fraction of the total T cell compartment (8-20%, depending on the cell subset), reinforcing the idea of a chronic and specific antigen stimulation. This widespread expansion supports sustained, antigen-driven T cell activation within mature ELS and aligns with the increased frequency of TEMRA T cells in mLM (**Fig. 5E**), reflecting the presence of antigen-experienced, clonally expanded T cells. In contrast to TCR repertoires, BCR repertoires in both iLM and mLM synovial samples exhibited similarly low numbers of unique clones, indicating that a small set of B cell clones dominates the repertoire (**Fig. 7D**). Clonal overlap analysis revealed higher sharing of BCR clones across patients, consistent with localized humoral responses within synovial ELS (**Fig. 7E**). Expanded B cell and plasma cell clones, defined by unique immunoglobulin sequences, were observed in the synovium, with variability across patients, reflecting heterogeneity in the humoral compartment of OA synovium (**Fig. 7F**). Whether such clonal expansions are specific to OA or also present in healthy synovium remains to be determined.

**Fig. 7.**
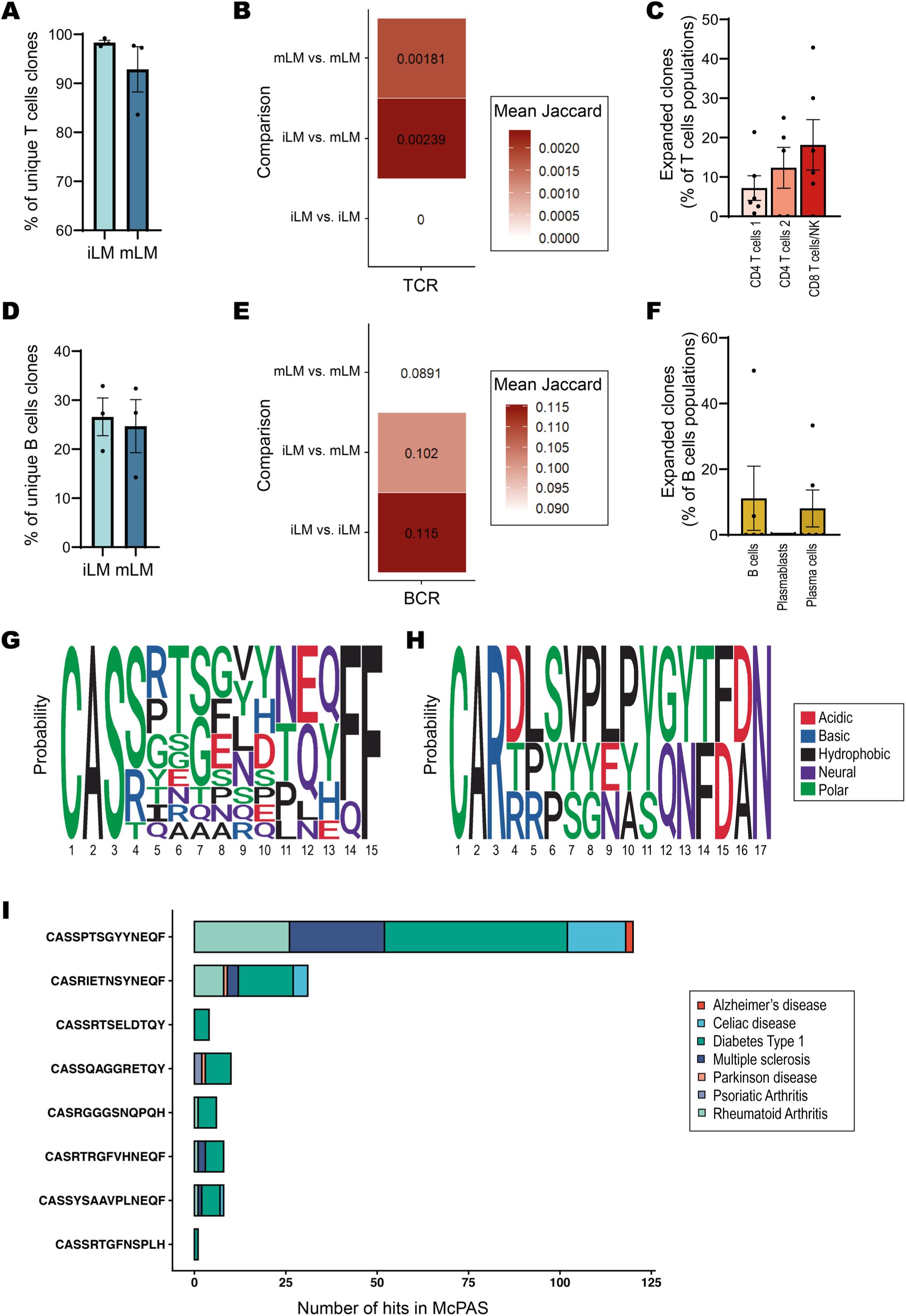
T and B cell repertoire analysis reveals antigen-driven expansion in mature OA ELS. **(A)** Percentage of unique T cell receptor (TCR) clones in immature lympho-myeloid (iLM) and mature lympho-myeloid (mLM) synovium. **(B)** Clonal overlap of TCR repertoires across patients (iLM vs. iLM; iLM vs. mLM and mLM vs. mLM), calculated using the Jaccard index. **(C)** Percentage of expanded T cell clones (clonal frequency >2) across CD4^+^ and CD8^+^ populations. **(D)** Percentage of unique B cell receptor (BCR) clones in iLM and mLM samples. **(E)** Clonal overlap of BCR repertoires across patients (iLM vs. iLM; iLM vs. mLM and mLM vs. mLM), calculated using the Jaccard index. **(F)** Percentage of expanded B cell clones (clonal frequency >2) across B cell, plasmablasts and plasma cells. **(G-H)** Sequence logos representing amino acid probabilities in TCR **(G)** and BCR **(H)** clonotypes. **(I)** Alignment of expanded TCR sequences against the McPAS-TCR database, showing the fraction of clones sharing ≥40% similarity with sequences previously associated with autoimmune diseases, such as type I diabetes, Alzheimer’s and Parkinson disease, celiac disease, multiple sclerosis, psoriatic and rheumatoid arthritis.

Similarity of amino-acid motifs among TCR (**Fig. 7G**) and BCR (**Fig. 7H**) clonotypes was observed. Notably, alignment of expanded TCR clones from OA synovium against the McPAS database^39^ showed that many shared ≥40% similarity with TCR previously associated with autoimmune diseases, predominantly type I diabetes, with fewer corresponding to RA, multiple sclerosis, celiac disease, and others (**Fig. 7I**). These data overall indicate that T cells within mature OA ELS (mLM) are antigen-experienced and may actively participate in local immune responses reminiscent of autoimmune conditions, whereas iLM samples display less clonal expansion and overlap, consistent with a less organized, immature immune microenvironment.

## DISCUSSION

A ‘one-size-fits-all’ therapeutic approach has been largely unsuccessful in OA, a highly heterogeneous disease, and the identification of synovial pathotypes represents a critical step toward patient stratification and precision medicine. Our findings demonstrate that these pathotypes, based on immune cell infiltration and distribution, are conserved across independent OA cohorts and are linked to joint tissue damage. This consistency underscores their potential as biomarkers for patient stratification and targeted therapeutic intervention. By defining immune endotypes, each characterized by distinct gene signatures, we move closer to a framework where therapies can be personalized to the molecular and cellular landscapes of the joint^2,7^.

Here, we notably highlighted that specific targets linked to joint remodeling (PI), angiogenesis and metabolism (DM), or leukocyte homing and activation (LM) could represent relevant candidates for pathotype-specific therapeutic interventions or biomarkers to guide precision medicine approaches. Among these candidates, Asporin (*ASPN*) and Osteoprotegerin (*TNFRSF11B)*, upregulated in PI synovium, are key regulators of extracellular matrix remodeling and bone turnover^40,41^, respectively, reflecting key pathways in joint homeostasis and OA progression. CD34 and AOC3 are two examples of angiogenesis-related molecules overexpressed in synovium exhibiting a DM pathotype. Targeting angiogenesis, which contributes to chronic inflammation and facilitates immune cell infiltration into the joint^42^, may be particularly relevant in this context. Moreover, we showed that metabolism-associated processes could also be associated with a DM pathotype. Given that synovial and infrapatellar fat pad form an anatomo-functional unit^43^, and that our study revealed a positive correlation between macrophage infiltration in both tissues, the potential interplay between these two compartments should be further investigated in OA, particularly in this DM pathotype. Indeed, synovium and infrapatellar fat pad are increasingly recognized as central players in OA progression^2,44^, and should not be overlooked when investigating disease mechanisms or developing targeted therapies. Finally, and as expected given the lymphoid infiltration that characterizes the LM pathotype, targeting immune checkpoint molecules (e.g., PD-1, CTLA-4), or chemokine receptors (e.g., CXCR4, CCR7) involved in lymphocyte recruitment and maturation, could represent a relevant strategy in this subgroup of patients. Notably, the LM pathotype is the most prevalent in end-stage OA, affecting approximately 50% of patients, highlighting the potential impact of such targeted approaches. As systemic biomarkers may also reflect whole joint pathology (e.g., cartilage degradation, synovial inflammation, metabolic perturbations)^8,10^, integrating synovial endotypes with circulating biomarkers in the future would also offer a promising avenue for non-invasive patient stratification.

Following the definition of disease endotypes using high-throughput transcriptomic analyses, digital spatial profiling revealed niche-specific transcriptomic profiles, highlighting the spatial heterogeneity of OA synovium. In line with the findings of Philpott et al.^45^, synovial lining and sublining compartments were found to possess distinct gene and protein signatures associated with their anatomical and functional identity. Moreover, we reported for the first time that each synovial niche displays pathotype-specific features, with PI lining and sublining enriched for innate immune signatures, DM lining showing stress-response and fibroblast activation programs, and LM lining and sublining characterized by adaptive immune activity and B/T cell-associated gene signature, underscoring the relevance of our stratification strategy. As previously demonstrated, synovial immune dysfunction in OA is associated with worse pain^45^, and future work should investigate whether specific pathotypes or niche organizations correlate with pain phenotypes, which could inform both mechanistic studies and clinical management.

While OA has historically been considered a non-autoimmune disease, emerging evidence suggests that adaptive immune responses may not just be bystanders during OA^2,18^, though their pathogenic role and clinical relevance remain less defined than in RA^46,47^. In line with these observations, the presence of ELS in the LM pathotype suggests a localized adaptive immune response. Here, for the first time, we describe the presence of mature ELS, characterized by the presence of CD21^+^CD23^+^ follicular dendritic cells, in about one-third of LM synovial tissues. This observation raises important questions about the functional role of these structures in OA and their potential as therapeutic targets. Importantly, our data indicate that the presence of mature ELS is associated with radiographic markers of joint damage, strongly suggesting that these structures may contribute to local tissue remodeling and disease severity, similarly to what has been observed in RA synovium^24^. These results are consistent with a recent report linking synovial lymphocyte and plasma cell infiltrates to greater structural damage^48^, together supporting a pathogenic role for mature ELS in OA. Our data also support the notion that the maturity of ELS is linked to cell density and could therefore serve as proxies for immune activity. The future translation of our findings into routine clinical practice will require robust and scalable methodologies. The automation of mLM detection using standard H&E staining combined with deep learning algorithms could facilitate its widespread adoption, allowing clinicians to rapidly assess the presence of mature ELS, better stratify patients, and potentially predict disease response, as already implemented in the field of oncology^49^.

Our integrated single-cell and spatial transcriptomic analyses reveal the structural organization and functional maturation of ELS in OA synovium. LM synovial tissues exhibiting mature ELS (mLM) were associated with a global rewiring of the synovial microenvironment, consistent with observations in other contexts where tertiary lymphoid structures shape local tissue organization and immune activity, such as in cancer^50^. We highlighted ligands such as CXCL8/3/2, CCL3/4, THBS1, and Cyclophilin A, along with their respective receptors, which coordinate immune-stromal interactions, lymphocyte recruitment, and tissue remodeling in mLM synovium, representing potential molecular targets for precision interventions. Notably, mLM synovium displays an increased proportion of TEMRA and naïve B cells, reflecting selective expansion of adaptive immune compartments and a functional reorganization of the immune landscape. For the first time, this study shed light on the structural organization of ELS in OA synovium, with CosMx spatial analyses providing unprecedented resolution of the cellular architecture and interactions between immune, stromal, and vascular components. We notably highlighted that mature OA ELS exhibited spatial organization similar to those observed in RA^36,51^, consistent with germinal center-like activity^47^. The presence of Tfh and GC B cells within mature OA ELS, consistent with patterns observed in RA, highlighted their functional maturity. Of note, although the RA tissues included in this study exhibited mature ELS and were from autoantibody-positive patients, they were collected during arthroplasty or synovectomy from patients treated with various disease modifying anti-rheumatic drugs, and not at the early stage of disease. This may have influenced the proportion of mature ELS observed in the RA synovial tissues analyzed herein, and the comparison with OA tissues. Indeed, the percentage of CD21^+^ RA synovial tissues can vary across cohorts, ranging from approximately 25% to over 70%, depending on disease activity and treatment regimen^46,52^. In addition, our study should now be extended to include synovial tissues from earlier stages of OA, in order to confirm the distribution of pathotypes, assess the presence of mature ELS from disease onset, and explore potential pathotype or ELS maturation shifts in longitudinal studies.

Our analysis of T and B cell repertoires in OA synovium provides novel insights into the functional maturation of ELS. In mLM, TCR repertoires exhibited a reduced percentage of unique clones, with increased clonal overlap across patients, indicative of local, antigen-driven T cell expansion. In contrast, BCR repertoires showed higher similarity across iLM and mLM patients. This pattern is reminiscent of tertiary lymphoid structures in autoimmune diseases and tumors, where T and B cell clonal dynamics are closely intertwined, and T cell activation can both precede and support B cell selection and plasma cell differentiation^47,53^. The predominance of expanded T cell clones in mature OA ELS supports the notion of an active and organized immune microenvironment that could contribute to local tissue remodeling and structural progression, consistent with the association between ELS maturity and radiographic joint damage. Notably, several of these expanded TCR sequences show similarity to clones previously described in other autoimmune diseases, reinforcing the concept of antigen-driven T cell selection within OA synovial ELS, potentially linked to the production of autoreactive antibodies reported in OA^20–22^. Although these repertoire findings were derived from a limited number of patients, calling for larger studies including patients at different stages of OA, as well as analyses of circulating T and B cell repertoires to validate and extend these observations, they underscore a unique immunological landscape in OA. Future studies could include antigen discovery approaches using MHC-peptide complexes to identify the stimuli driving T and B cell clonal expansion.

Overall, our study provides a comprehensive atlas of OA synovial pathotypes/endotypes, revealing how immune cell composition, spatial organization, and adaptive immune repertoires converge to define the functional maturation of ELS. Importantly, these findings extend beyond descriptive pathology: they provide actionable insight for patient stratification, with potential to guide targeted therapies and precision medicine approaches. These observations underscore the existence of distinct immunological endotypes in OA, setting the stage for interventions tailored to the molecular and cellular landscape of individual patients, and advancing precision medicine in OA.

## MATERIALS AND METHODS

### Patients

Patients included in Cohort 1 have been described previously^13^, and were enrolled in the EMR Biobank study [REC Ref. No. 07/Q0605/29] (London, UK). Cohort 2 is composed of 132 patients with confirmed diagnosis of knee OA, according to EULAR recommendations^54^, enrolled at Nantes University Hospital [DC-2017-2987] (Nantes, France). OA patients underwent total joint replacement and were excluded if they were diagnosed with other joint diseases. Clinical data, including age, sex, BMI (cohorts 1 and 2) and time since diagnosis (cohort 2) were collected for each patient. Synovial tissues were collected (cohorts 1 and 2), as well as matched cartilage from the tibial plateau and infrapatellar fat pad when available (cohort 2) (**Table. S1**). Radiological severity of patients exhibiting a LM pathotype was assessed by two independent readers experienced in the radiographic evaluation of OA, according to the following criteria: joint space narrowing, osteophytes, subchondral sclerosis and cysts, condyle impaction and deformation, presence of mono, bi, or tricompartimental OA^55,56^.

Synovial tissues from RA patients diagnosed according to the European Alliance of Associations for Rheumatology (EULAR) criteria^57^ were obtained during arthroplasty or synovectomy. Patients had established disease, were treated with disease-modifying anti-rheumatic drugs (DMARDs) and were enrolled at Nantes University Hospital under the approval DC-2011-1399. All patients had active disease (DAS28 > 2.6) and were seropositive for anti-citrullinated protein antibodies and/or rheumatoid factor.

### Sample collection and processing

Upon collection, OA synovial and fat pad tissues were divided into two similar pieces, one cryopreserved in Cryostor CS10 (Sigma-Aldrich, C2874) according to previously described procedures^58^ for transcriptomic analyses or spectral flow cytometry, the other fixed in 4% paraformaldehyde (PFA) for 24 hours at 4°C and embedded in paraffin for histology. RA synovial tissues were fixed in 4% PFA for 24 hours at 4°C upon collection and embedded in paraffin for histology. Tibial plateaus were fixed in 4% PFA for a week at 4°C and divided into 8 pieces using a Dremel saw (Dremel) before decalcification in a solution of ethylenediaminetetraacetic acid (EDTA) at 0.636M, pH 7.5-8, at 4°C for several weeks until complete decalcification. Cartilage samples were then embedded in paraffin for histology (FFPE).

### Histology

3µm-thick synovial tissue FFPE sections were stained with hematoxylin and eosin (H&E) to check tissue quality and assign Krenn scores^28^ (0-1, no synovitis; 2-4, low-grade synovitis; 5-9, high-grade synovitis). As previously described^13^, sections underwent IHC staining for T cells (CD3, M7254 Agilent Technologies), B cells (CD20, M0755, Agilent Technologies), macrophages (CD68, M0814, Agilent Technologies) and plasma cells (CD138, M7228 Agilent Technologies) to determine synovial pathotypes (pauci-immune, diffuse-myeloid and lympho-myeloid). CD21 (M0784, Agilent Technologies) and CD23 (ab92495, Abcam) staining were also performed on lympho-myeloid tissues to determine the maturity of lymphoid aggregates. Fat pad 3 µm-thick sections were stained with H&E to assess the degree of fibrosis, vascularization, and lymphocytic infiltration, using a semi-quantitative scoring system developed by our team based on histopathological features commonly evaluated in adipose tissues (each criterion scored on a 0-4 scale). Sections also underwent immunohistochemical staining to assess macrophage infiltration (CD68). The characterization of cartilage degradation was performed using the Osteoarthritis Research Society International (OARSI) score^29^ on 4µm-thick sections stained with Safranin-O Fast-green. All scoring were performed by two independent observers.

All sections were digitally scanned using Nanozoomer S360 (Hamamatsu Photonics).

### RNA extraction and bulk RNA sequencing

RNA was extracted using a Trizol/Chloroform method (Cohort 1, 6 PI, 38 DM, 35 LM) or using Nucleospin columns (Macherey-Nagel; Cohort 2, 3 PI, 6 DM and 6 LM). RNA library preparations and sequencing reactions were conducted at GENEWIZ, LLC. (South Plainfield, NJ, USA; Cohort 1) and at GenoToul (Toulouse, France; Cohort 2). The libraries were sequenced on the Illumina NovaSeq 6000 instrument using a 2x150 Paired End configuration. The fastq files were trimmed using the Fastp function. Indexation and alignment were then performed using the index and quant functions, respectively, in SALMON (ensembl release 113 homo sapiens). Batch adjustment was performed using CombatSeq from the sva package (version 3.50.0). Analyses were performed under R (version 4.3.3), genes with less than 3 counts per million (CPM) and identified in fewer than 4 samples were filtered out, differentially expressed genes (DEG) were obtained using the DESeq2 package (version 1.42.1). Gene ontology (GO) analyses were performed using the Enrichr package (version 3.2) using “GO_Molecular_Function_2021”, “GO_Cellular_Component_2021” and “GO_Biological_Process_2021” databases. Single-sample Gene Set Enrichment Analysis (ssGSEA) was performed using the GSVA package (version 2.2.0) to estimate the enrichment scores of predefined gene sets for each individual sample. Enrichment scores were calculated by ranking genes within each sample and computing a normalized enrichment score for each gene set. Cell-specific gene sets were derived from FANTOM5 data to enable relative quantification of cell populations in bulk RNA-seq and spatial GeoMx datasets. Module scores for each cell type were calculated using the AddModuleScore function from the Seurat R package^59–61^.

### GeoMx Digital Spatial Profiler (DSP) whole transcriptome atlas (WTA) and protein spatial analyses

3-µm FFPE synovial tissues sections were prepared from 16 OA (4 PI, 6 DM and 6 LM) and 4 RA (all LM) patients were profiled using the NanoString GeoMx DSP Whole Transcriptome Atlas (WTA) assay, as previously described^33,34^, and the “Immune cell profiling”, “Immune cell typing” and “Immune cell activation status” protein panels (total of 35 protein analyzed). The following fluorescent markers were used to determine the morphology of the tissue: CD68-AF647 for macrophages (NBP2-34587AF647, Novus Biologicals), CD20-AF594 for B cells (NBP2-47840AF594, Novus Biologicals), and CD3-AF532 (uncoupled CD3 antibody A0452 Dako, and secondary antibody anti rabbit AF532, A-11009, ThermoFisher) for T cells; nuclei were counterstained with Syto13 (S7575, ThermoFisher). In situ hybridizations with the NanoString GeoMx WTA panel, including 18,677 gene probes, and GeoMx fluorescent antibodies were performed on serial sections, according to the manufacturer’s instructions (Nanostring, MAN-10150-01 and MAN-10089-08). Freeform polygon-shaped regions of interest (ROIs) containing a minimum of 200 cells were selected on immunofluorescent-stained images to include synovial tissue 1/ lining (characterized by the presence of CD68^+^ cells); 2/ sublining (characterized by the absence of CD3^+^ and CD20^+^ cells aggregates); 3/ ectopic lymphoid structures or ELS (characterized by the presence of CD3^+^ and CD20^+^ cells forming aggregates). Subsequently, individual segmented areas within ROIs were photocleaved by the NanoString GeoMx DSP and oligonucleotides collected into separate wells of two 96-well collection plates for RNA and protein analyses. The dataset generated included 184 (RNA) and 159 (Protein) ROIs from 24 patients (before QC). NTC water wells were used for quality control checks.

#### Protein

Collected samples were processed using the NanoString nCounter® according to the manufacturer’s instructions (Nanostring MAN-10089-08), generating proteomic data for each ROI. Quality control and normalization were performed onto the instrument according to the manufacturer’s instructions (Nanostring, MAN-10154-01). 154 ROIs passed the QC step for further proteomic analyses. Subsequent analyses were conducted in R (version 4.3.3)

QC and normalization were performed using the GeoMx DSP instrument. Differentially expressed proteins (DEP) analysis was performed using the DESeq2 package (version 1.42.1). Gene set enrichment analysis (GSEA) was performed using the C5 ontology genesets database using GSEA software (Broad Institute, version 4.4.3).

#### WTA

NanoString GeoMx WTA sequencing reads were compiled into FASTQ files corresponding to each ROI. FASTQ files were converted to digital count conversion files using the NanoString GeoMx NGS DnD Pipeline. The GeoMxTools package (version 3.6.2) was used for QC and normalization (quant method) starting from Nanostring DCC and PKC files generated from the GeoMx DSP instrument. ROIs with a gene detection rate of less than 4% and genes detected in less than 10% of the ROIs were excluded. 137 ROIs passed the QC step for further transcriptomic analyses. DEG were obtained using the mixedModelDE function from the GeomxTools package. 3D volcano plots were generated using ‘volcano3D’ package version 2.0.9.

### scRNA-seq and TCR/BCR sequencing

#### Synovial tissue digestion

Cryopreserved synovial tissues were thawed and rinsed twice in RPMI medium supplemented with 10% fetal bovine serum (FBS) at room temperature (10 and 5 min, respectively). Tissues were then subjected to a final wash in RPMI alone for 5 min. Tissues were digested both enzymatically and mechanically using gentleMACS™ C tubes (Miltenyi Biotec) in a digestion medium consisting of 400 µg/mL DL Liberase (Roche, 5466202001) and 80 µg/mL DNase I (Sigma-Aldrich, 10104159001) prepared in RPMI. Dissociation was performed using the “37C_mTDK2” program on the gentleMACS™ Octo Dissociator (Miltenyi Biotec). Enzymatic digestion was stopped by adding RPMI supplemented with 10% FBS, and the resulting cell suspension was filtered through a 70-µm cell strainer. Cells were then centrifuged at 330g for 6 min at 4 °C, and the cell pellets were resuspended in RPMI medium containing 10% FBS. Prior to scRNA-seq, the cell suspension was enriched for CD45^+^ cells using a Dead Cell Removal Kit (Miltenyi Biotec, 130-090-101) followed by positive selection with human CD45 MicroBeads (Miltenyi Biotec, 130-045-801), according to the manufacturer’s instructions.

#### scRNA-seq libraries preparation

Single-cell RNA-seq was performed using the BD Rhapsody™ Single-Cell Analysis System, the libraries were generated following the manufacturer’s instructions. Each single-cell suspension was labeled for sample multiplexing using oligonucleotide-conjugated sample tags (Single Cell Sample Multiplexing Kit, 633781). Cells were then captured on BD Rhapsody microwell cartridges. Four libraries were prepared from the same captured cells: whole transcriptome analysis (WTA, BD Rhapsody WTA Amplification Kit, 633801) libraries, TCR and BCR (TCR/BCR Next Amplification Kit, 667058) libraries, and sample multiplexing libraries. Following reverse transcription and cDNA amplification (Rhapsody cDNA Kit, 633773), libraries were constructed according to BD protocols, quality-controlled, and sequenced on an Illumina NextSeq2000 platform according to the manufacturer’s recommendations.

### CosMx Spatial Molecular Imager (SMI) analyses

3-µm FFPE synovial tissues sections were prepared from 2 OA and 1 RA patients and profiled using the NanoString CosMx Spatial Molecular Imager (SMI) Human 6000 Gene Panel. 49 fields of view (FOV) were selected across the tissue sections (including a total of 10 ELS FOV). Single-cell resolution transcript counts were processed and quality-controlled using the CosMx software pipeline and subsequently analyzed, as described in the next section.

### scRNA-seq and CosMx spatial transcriptomic integration and analyses

Integration of scRNA-seq and CosMx spatial transcriptomics data was performed using Seurat (version 5.4.0) in R. To ensure comparability across platforms, genes detected in both datasets were identified, resulting in a set of 6,044 common genes, which were retained for all subsequent analyses. New Seurat objects were generated for scRNA-seq and CosMx data using only these shared genes, while preserving the original metadata. QC were performed based on nCount, nFeatures and cell size (for CosMx only), both datasets were independently normalized using SCTransform. 6 090 cells were analyzed in the scRNA-seq dataset and 115 864 in the CosMx dataset following QC. Both objects were merged, and dimensionality reduction was performed using principal component analysis, followed by batch correction and data harmonization using Harmony (version 1.2.4), accounting for sample-specific effects. Multiple values of the Harmony diversity clustering penalty (theta) were evaluated, and the optimal parameter was selected based on improved mixing of datasets and preservation of biological structure (theta 2). Following Harmony integration, cell clustering (resolution 0.7), and visualization were performed, and two-dimensional representations were generated using UMAP. Cell proportions are compared using ‘speckle’ package v1.8.0. For spatial analyses of ELS, a free-form selection tool was used to delineate ELS regions, enabling precise attribution of individual cells to either “Within ELS” or “Outside ELS” areas for spatial characterization (SeuratSpatialSelector package v0.1.1 github.com/RomainGuiho/SeuratSpatialSelector based on plotly package version 4.11.0). Neighborhood analyses were performed using CatsCradleSpatial (github.com/AnnaLaddach/CatsCradle, version 1.2.0) following extended Delaunay triangulation of combinatorial radius 4. Interactions are visualized using visNetwork package v2.1.4.

### TCR and BCR sequencing analyses

TCR and BCR V(D)J sequencing data generated using the BD Rhapsody platform were processed in R using the scRepertoire package^62^. For repertoire-level analyses, TCR and BCR data were aggregated per sample and stratified according to patient group (iLM versus mLM). The proportion of unique clonotypes was calculated to assess repertoire diversity. Clonal expansion was defined as clonotypes detected in more than two cells. Clonal overlap between patients and their maturity status was quantified using the Jaccard index. Single-cell resolved analyses were performed by integrating TCR and BCR information into the scRNA-seq dataset after removal of V(D)J genes from the expression matrix to avoid clustering bias. Clonal expansion was assessed across major T cell (CD4^+^ and CD8^+^) and B cell populations (B cells, plasmablasts, and plasma cells). Amino acid sequence features of expanded TCR and BCR clonotypes were visualized using sequence logo representations generated with the ggseqlogo R package^63^. Expanded TCR amino acid sequences were aligned against the McPAS-TCR database^39^, and the proportion of clonotypes sharing ≥40% sequence similarity with previously reported antigen-associated TCRs in autoimmune and immune-mediated diseases was calculated.

### Spectral flow cytometry

Following tissue digestion as described above for scRNA-seq, cells were processed for flow cytometry staining. All antibodies used are listed in **Table. S2**, and a detailed staining procedure is provided in **Table. S3**. Briefly, after viability staining, cells were washed and subjected to sequential surface marker staining. Cells were then fixed with 2% PFA and stored in 1x Tandem Stabilizer buffer (BioLegend, 421802) until data acquisition on a 5 lasers Cytek Aurora spectral flow cytometer. Flow cytometry data were analyzed using OMIQ software (Dotmatics). Quality control was performed using the PeacoQC algorithm on live, singlet CD45^+^ cells.

### Statistics

Prior to statistical analysis related to clinical and histopathological data, missing values were imputed using the median of the corresponding variable, in order to minimize the influence of outliers and preserve the overall data distribution. The riverplot presented in Figure 1 was generated with SankeyMatic (https://sankeymatic.com/). PCA, PLS-DA, bubble plots and associated analyses were generated using RStudio (version 2025.05.1).

All other statistical analyses were performed using GraphPad Prism-v9 software (GraphPad, San Diego, CA, USA). p-values<0.05 were considered significant, and each test used is specified in figures and table legends.

## ACKNOWLEDGEMENTS

The authors thank the GeT-Santé facility (I2MC, Inserm, Génome et Transcriptome, GenoToul, Toulouse, France) for the advice and technical contribution to the generation of bulk RNA-seq data (cohort 2) (Emeline Lhuillier). The authors thank the « Plateforme de recherche en oncologie des hospices civils de Lyon », as well as the Nanostring technical support team (Ana Ortalli) for the GeoMx experiment. We acknowledge the MicroPICell core facility (SFR Bonamy, BioCore, Inserm UMS 016, CNRS UAR 3556, Nantes, France), member of the Scientific Interest Group (GIS) Biogenouest, IBISA, and the national infrastructure France-Bioimaging supported by the French national research agency (ANR-24-INBS-0005 FBI BIOGEN. We also acknowledge the SC3M facility from the Inserm/Nantes Université/ONIRIS UMR1229 RMeS Laboratory and SFR Bonamy for their help in histology and imaging, and the Inserm/Nantes Université/ONIRIS UMR1229 RMeS SC4BIO facility for their help in processing tissues and RNA extractions. We thank the GENOMAX core facility (Strasbourg, France) for sequencing. Finally, the authors would like to sincerely thank the patients who generously participated in this study, as well as the clinical team and surgeons from the Orthopedic Department of Nantes University Hospital for providing patients’ samples, and the clinical fellows from the Rheumatology Department for their help in retrieving clinical data.

## AUTHOR CONTRIBUTIONS

Conceptualization: C.P., M.-A.B., J.G.

Methodology: N.G., A.C., R.G., C.C., A.O., C.L., J.B., J.D.L., L.F.J., J.P., N.B., R.D., L.D., S.B., N.D., S.P., D.W., B.L.G., M.L., D.M.

Investigation: N.G., A.C., R.G., C.C., A.O., C.L., J.B., J.D.L., L.F.J., M.A., M.L.M., A.D., L.D., D.W., B.L.G., M.L., F.B., A.N., D.M.

Visualization: N.G., A.C., R.G., C.C., N.D., M.L.

Funding acquisition: C.V., M.-A.B., J.G., M.L., C.P.

Supervision: M.-A.B., J.G.

Writing - original draft: N.G., A.C., R.G., M.-A.B.

Writing - review & editing: All authors read, critically revised, and approved the final manuscript for publication.

