## Supplementary figures and tables for "Molecular and spatial profiling identifies immune endotypes for the stratification of OA patients"

- S1
- S2
- S3
- S4
- S5
- S6
- S7
- S8
- S9
- S10

### Supplementary Tables

- S1
- S2
- S3

**Supplementary Figure S1**

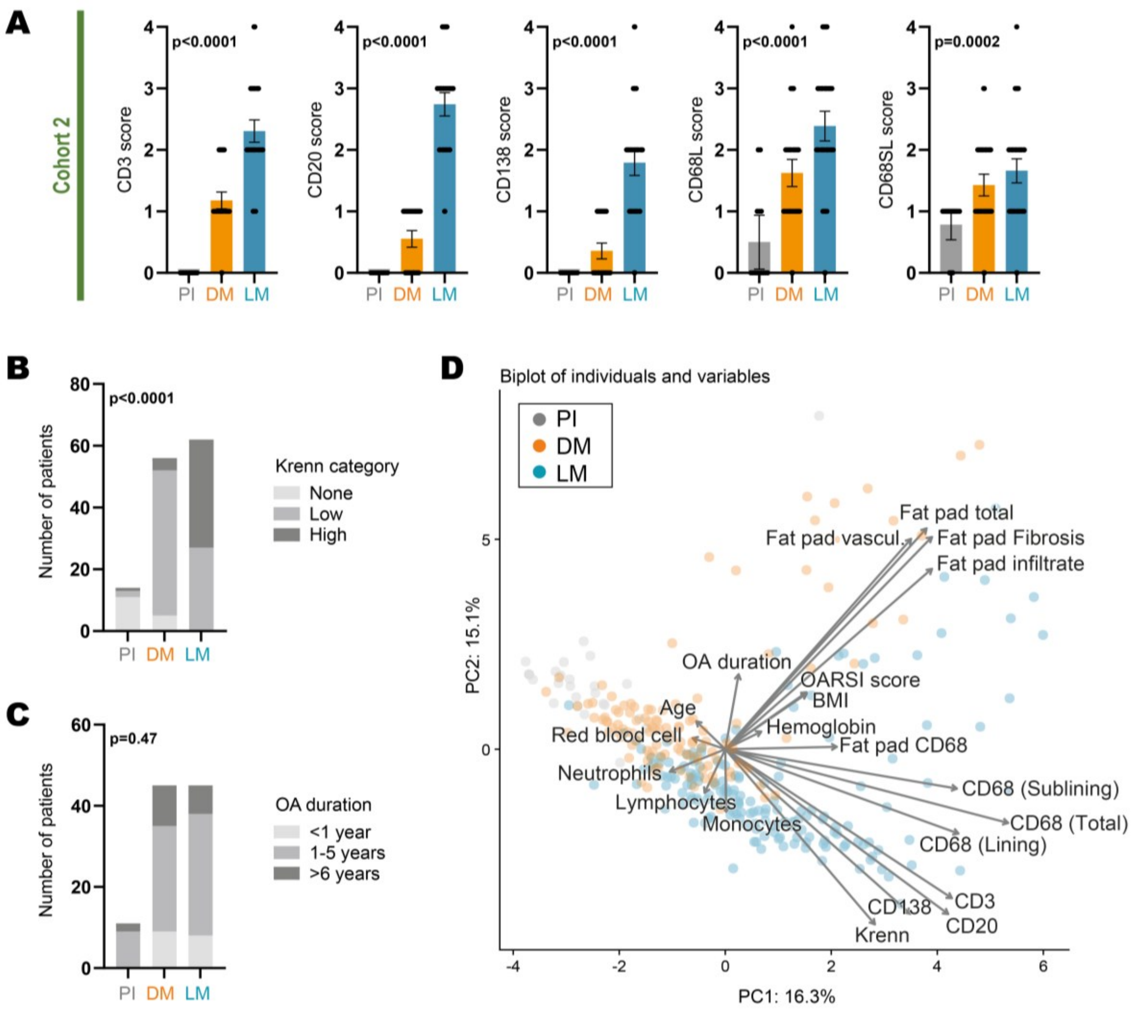


**Fig. S1.** ***Associated with main Figure 1.*** **(A)** CD3, CD20, CD138, lining CD68 (CD68L), and sublining CD68 (CD68SL) semi-quantitative scores in synovial tissues from OA patients belonging to Cohort 2, presenting a pauci-immune (PI, n = 14,), diffuse-myeloid (DM, n = 56), or lympho-myeloid (LM, n = 62). p-values were assessed by the Kruskal-Wallis test and Dunn's post-test, comparing PI, DM and LM groups for each marker. Individual values, mean and SEM are shown. **(B)** Bar chart representing the distribution of Krenn categories ("none" 0–1, "low" 5–4, and "high" 5–9) in each pathotype group PI, DM and LM) in synovium from OA belonging to Cohort 2 (n = 132). p-value was calculated using the Fisher's exact test. **(C)** Bar chart representing the distribution of time since diagnosis (OA duration, less than a year, between 1 and 5 years, and over 6 years in each pathotype group PI, DM and LM) in synovium from OA belonging to Cohort 2 (n = 132). p-value was calculated using the Fisher's exact test. **(D)** PCA biplot, the length and direction of variable vectors indicate their influence to PC1 and PC2, which account for 16,5% and 15,1% of variance, respectively. BMI, Body Mass Index; CD, Cluster of Differentiation; OARSI, Osteoarthritis Research Society International.

### Supplementary Figure S2

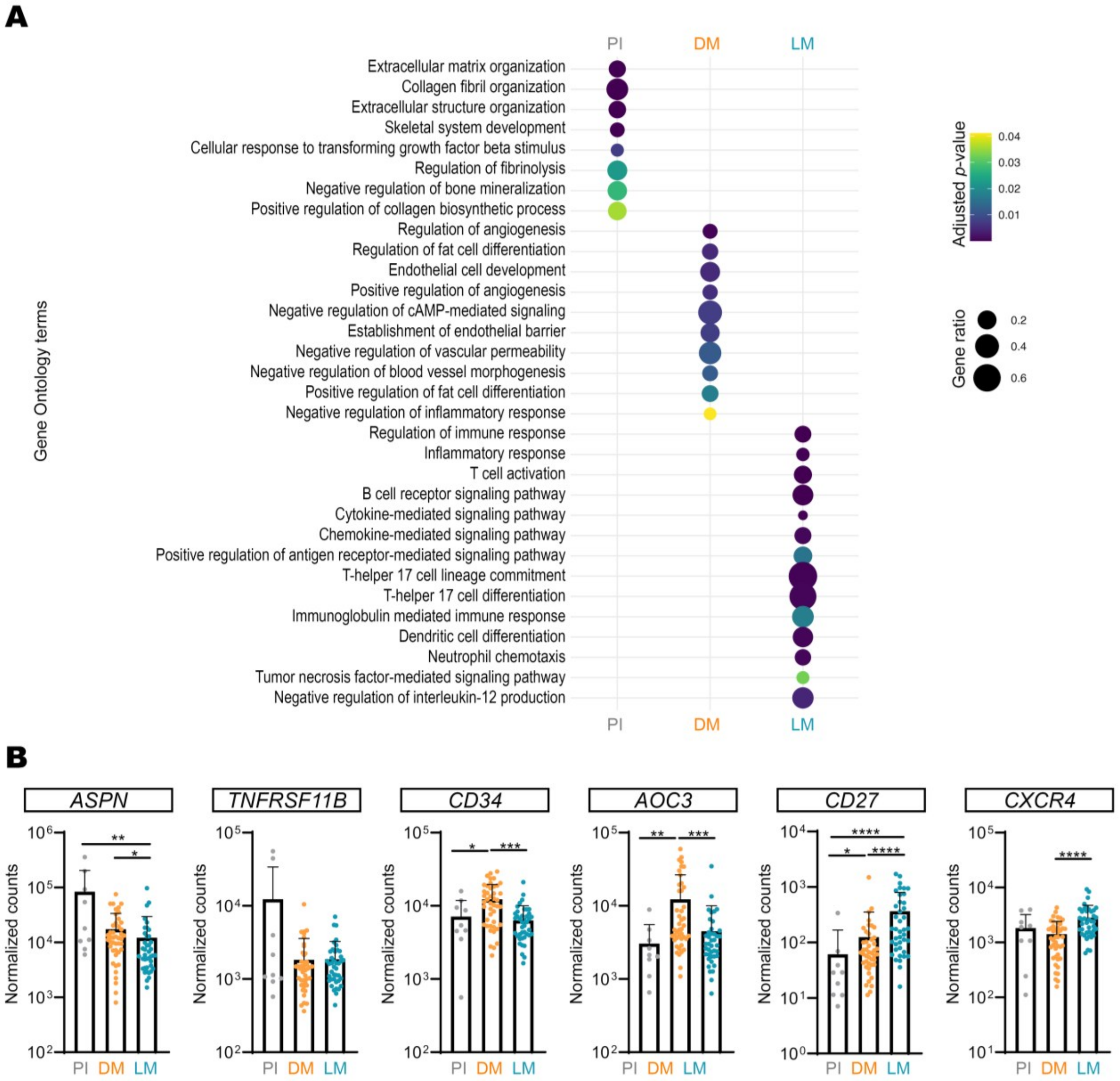


**Fig. S2.** ***Associated with main Figure 2.*** **(A)** Bubble plot representing selected enriched Gene Ontology (GO) terms associated with activated biological pathways in each of the three pathotypes (pauci-immune PI, diffuse-myeloid DM, and lympho-myeloid LM). The size of each bubble reflects the gene ratio (*i.e.,* the proportion of genes identified in the analysis relative to the total number of genes annotated to the GO term), while the color code indicates the statistical significance (adjusted p-value). **(B)** Histograms presenting the expression levels (normalized counts) of selected genes of interest for each pathotype (PI in grey, DM in orange and LM in blue), assessed by bulk RNA sequencing. Individual values, mean and SD are shown, p-values were calculated using the Kruskal-Wallis test with Dunn's post-test, * p<0.05; ** p<0.01; *** p<0.001; **** p<0.0001. ASPN, Asporin; TNFRSF11B, TNF Receptor Superfamily Member 11b or osteoprotegerin; CD34 and CD27, Cluster of Differentiation 34 and 27; AOC3, Amine Oxidase Copper Containing 3; CXCR4, C-X-C chemokine receptor type 4.

**Supplementary Figure S3**

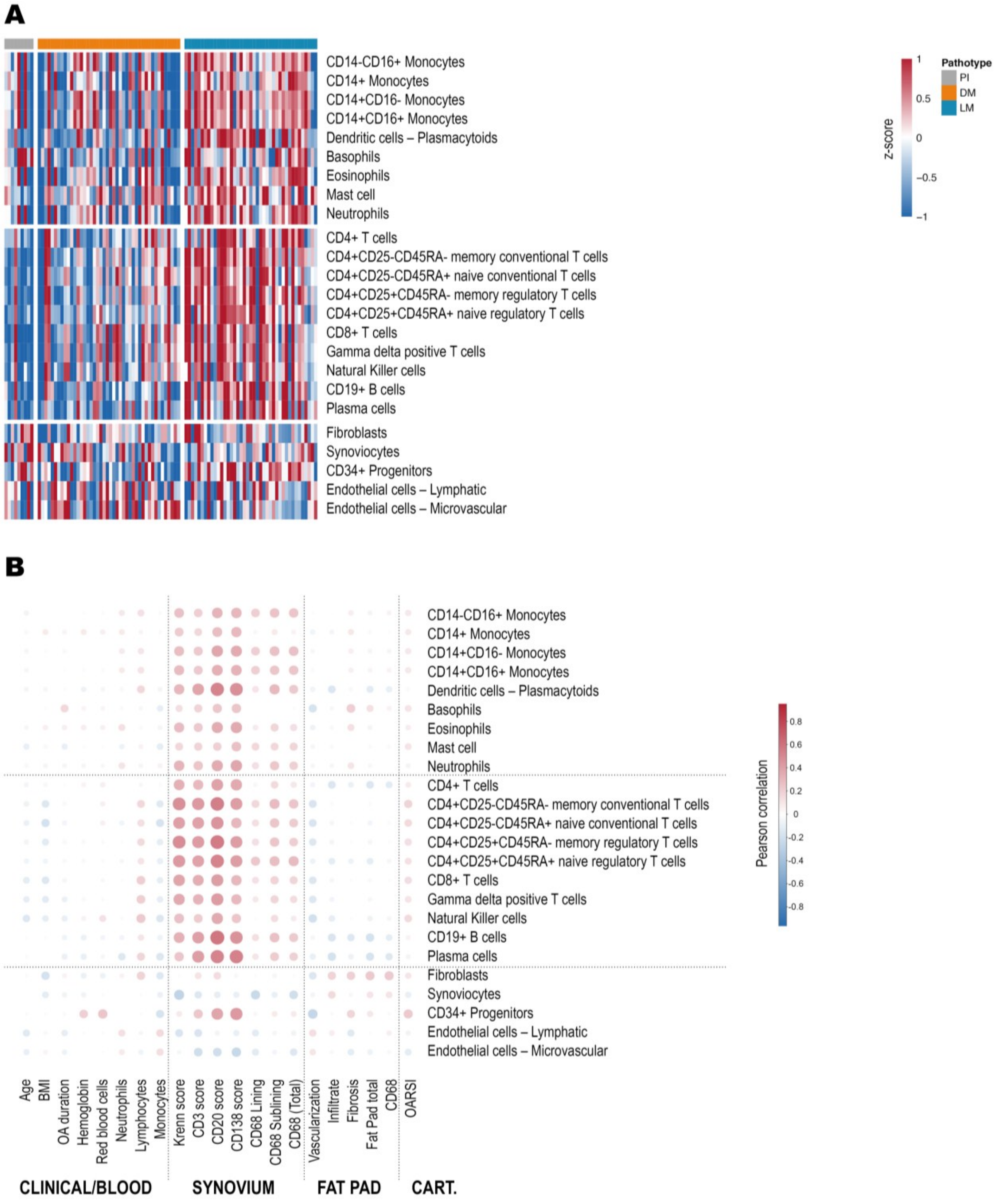


**Fig. S3.** ***Associated with main Figure 2.*** **(A)** Cell-specific gene set scores (derived from FANTOM5, as described in the Materials and Methods section) were used for the relative quantification of cell populations in synovial samples analyzed by bulk RNA sequencing. The blue/red scale indicates the module score for each population. **(B)** Correlation heatmap presenting Pearson correlation of cell-specific modules against clinical and histological data. The blue/red scale indicates the correlation coefficient. BMI, Body Mass Index; CD, Cluster of Differentiation; OARSI, Osteoarthritis Research Society International.

**Supplementary Figure S4**

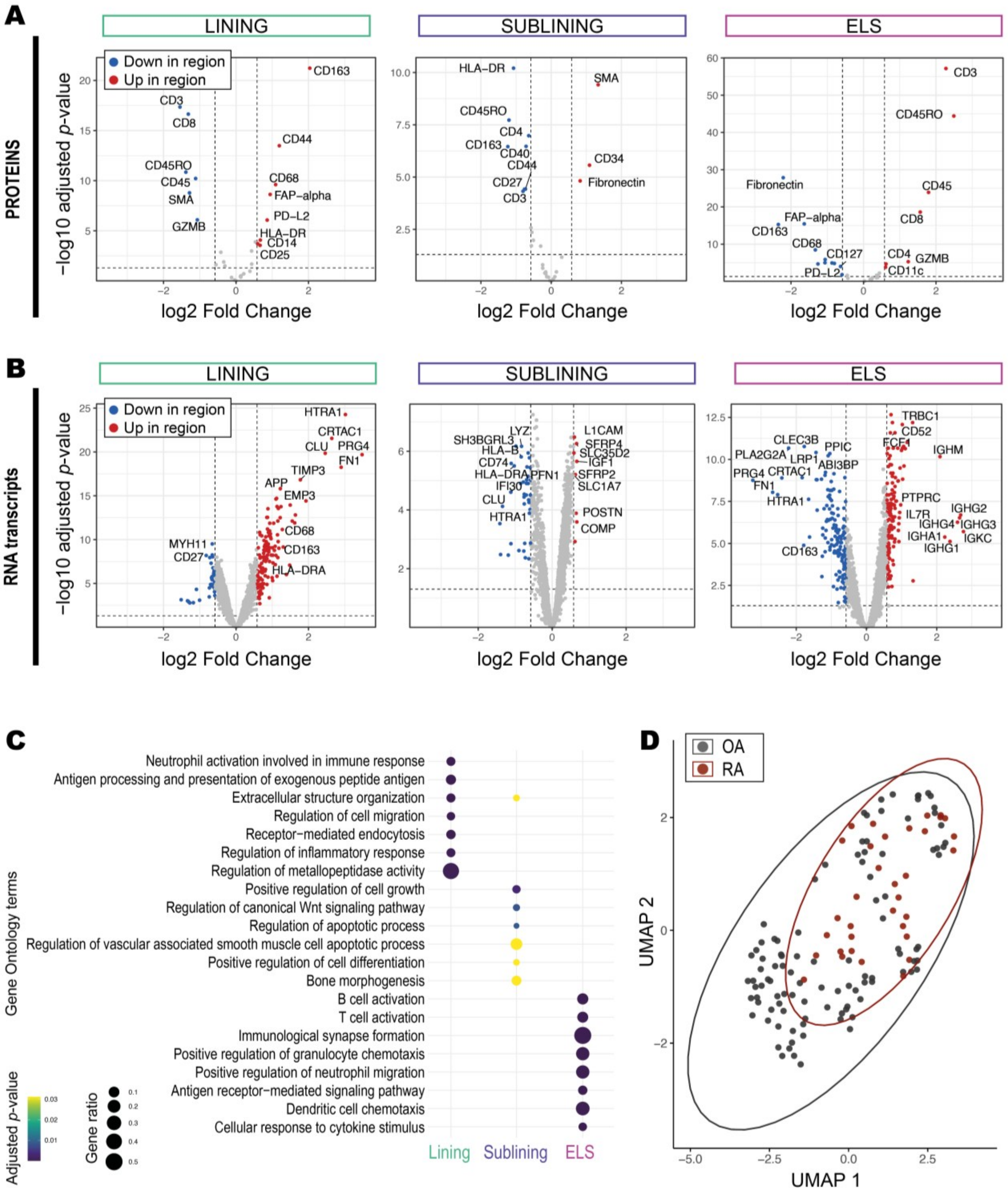


**Fig. S4. *Associated with main Figure 3.* (A, B)** Volcano plots presenting differentially expressed proteins **(A)** and genes **(B)** for each region of interest (ROI, lining, sublining and ectopic lymphoid structure or ELS) assessed by GeoMx Digital Spatial Imager (DSP). Each dot represents a protein **(A)** or gene **(B)**, the x-axis indicates the –log10(adjusted p-value) and the y-axis indicates the log2(fold-change) in expression between ROIs. Proteins **(A)** or genes **(B)** significantly enriched or downregulated in each ROI are highlighted in red and blue, respectively. **(C)** Bubble plot representing selected enriched Gene Ontology (GO) terms associated with activated biological pathways in each of the three ROIs (lining, sublining and ELS). The size of each bubble reflects the gene ratio (i.e., the proportion of genes identified in the analysis relative to the total number of genes annotated to the GO term), while the color code indicates the statistical significance (adjusted p-value). **(D)** Uniform Manifold Approximation and Projection (UMAP) projection of all ROIs analyzed using the GeoMx DSP from osteoarthritis (OA, black) and rheumatoid arthritis (RA, red) synovial tissues. Each dot represents an individual ROI, colored by its tissue of origin, as indicated in the legend.

**Supplementary Figure S5**

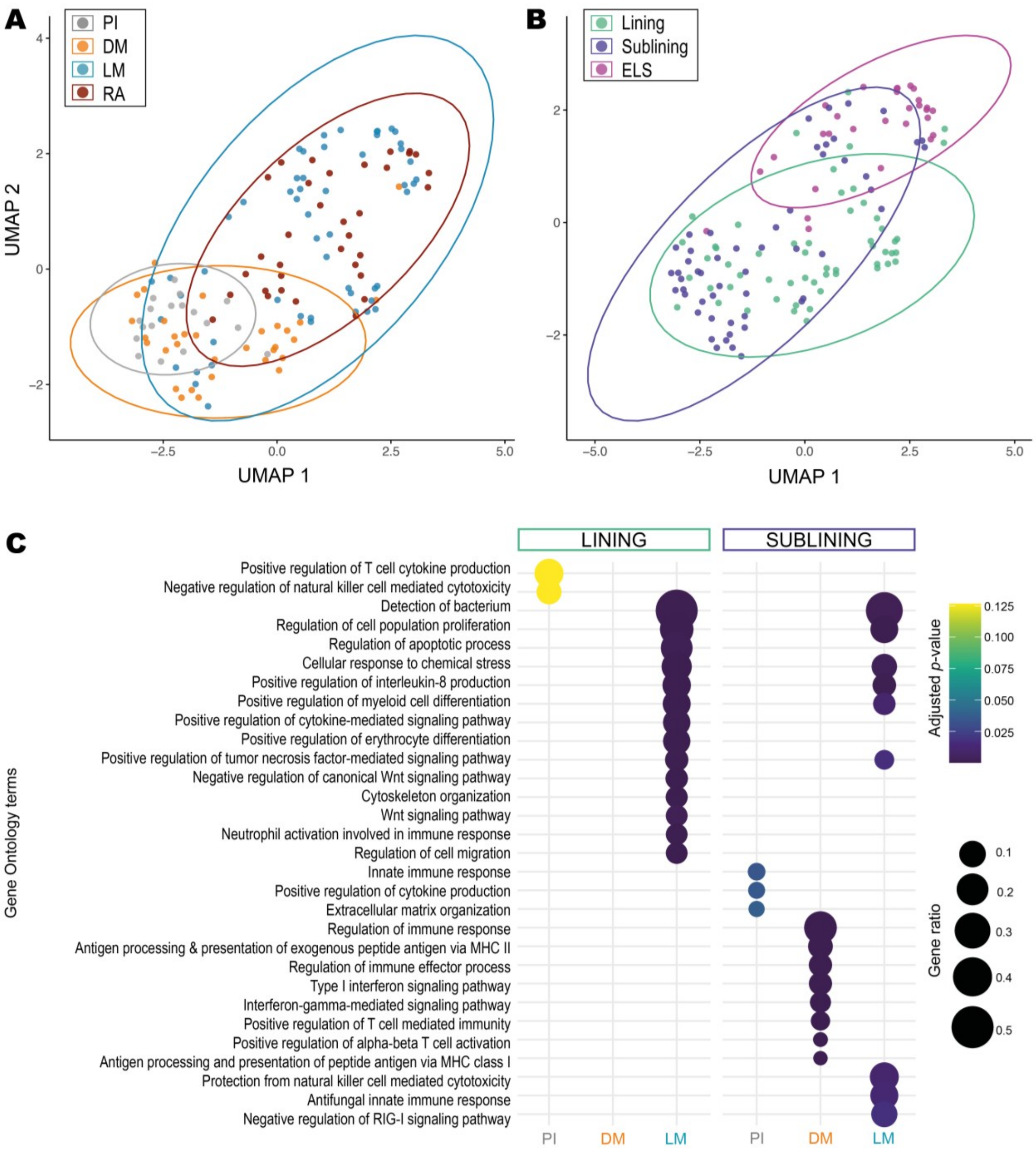


**Fig. S5.** ***Associated with main Figure 3.*** **(A-B)** Uniform Manifold Approximation and Projection (UMAP) projection of all region of interest (ROIs) analyzed using the GeoMx DSP from pauci-immune PI, diffuse-myeloid DM, lympho-myeloid LM, and rheumatoid arthritis RA tissues **(A)** or of each region of interest, namely lining, sublining and ectopic lymphoid structures (ELS) **(B)**. Each dot represents an individual ROI, colored by its tissue of origin, as indicated in the legend. **(C)** Bubble plot representing selected enriched Gene Ontology (GO) terms associated with activated biological pathways in each of the three pathotypes (pauci-immune PI, diffuse-myeloid DM, and lympho-myeloid LM) for lining and sublining ROIs. The size of each bubble reflects the gene ratio (i.e., the proportion of genes identified in the analysis relative to the total number of genes annotated to the GO term), while the color indicates the statistical significance (adjusted p-value).

**Supplementary Figure S6**

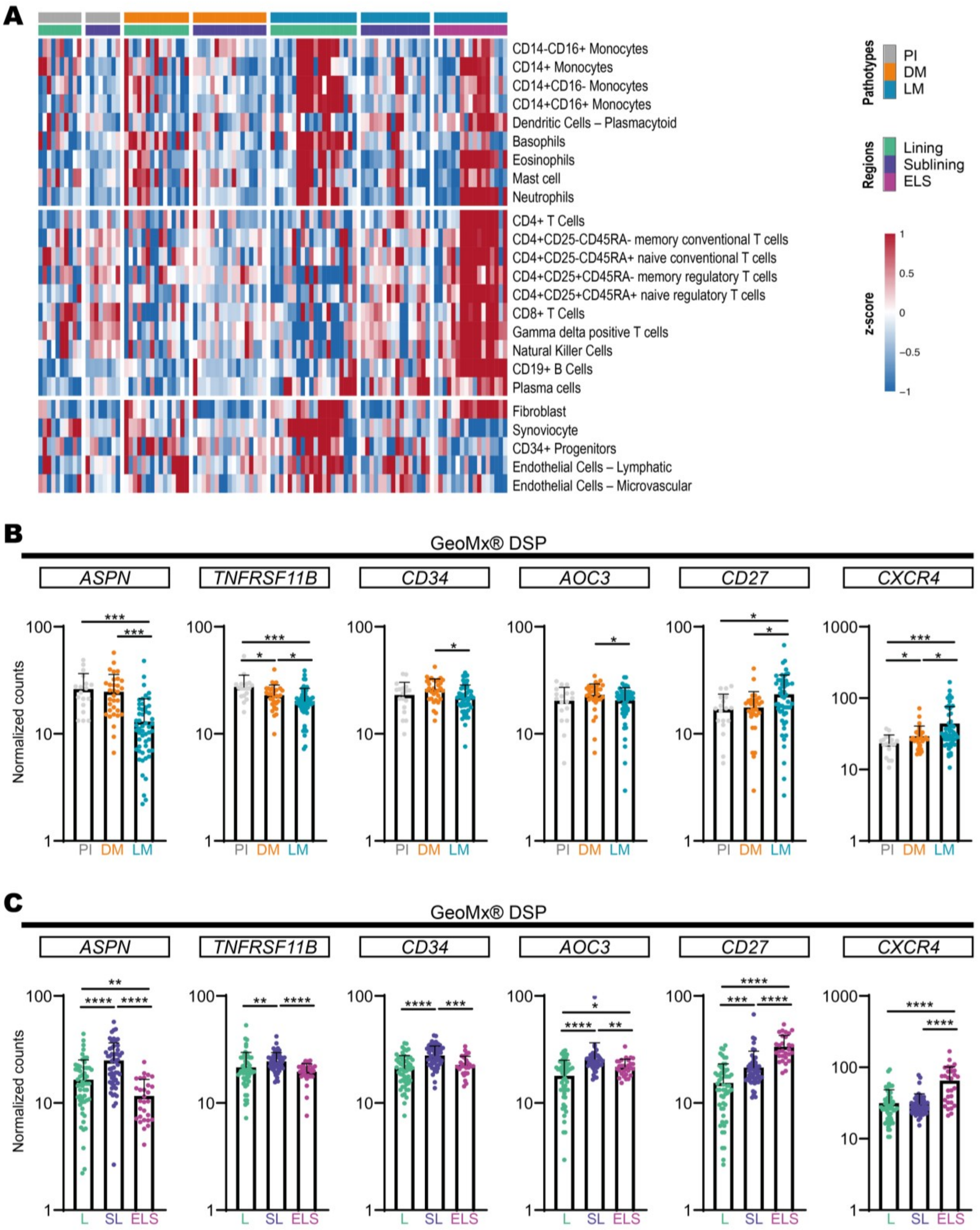


**Fig. S6.** ***Associated with main Figure 3.*** **(A)** Cell-specific gene set scores (derived from FANTOM5, as described in the Materials and Methods section) were used for the relative quantification of cell populations in synovial samples analyzed by GeoMx Digital Spatial Profiler. The blue/red scale indicates the module score for each population. PI, pauci-immune PI; DM, diffuse-myeloid; LM, lympho-myeloid. **(B)** Histograms presenting the expression levels (normalized counts) of selected genes of interest for each pathotype (PI in grey, DM in orange and LM in blue), assessed by GeoMx Digital Spatial Profiler (DSP). **(C)** Histograms presenting the expression levels (normalized counts) of selected genes of interest in each synovial niche (L, lining in green; SL, sublinining in purple; ELS, ectopic lymphoid structure in pink), assessed by GeoMx DSP. (**B-C**) Individual values, mean and SD are shown, p-values were calculated using the Kruskal-Wallis test with Dunn's post-test, * p<0.05; ** p<0.01; *** p<0.001; **** p<0.0001. ASPN, Asporin; TNFRSF11B,

TNF Receptor Superfamily Member 11b or osteoprotegerin; CD34 and CD27, Cluster of Differentiation 34 and 27; AOC3, Amine Oxidase Copper Containing 3; CXCR4, C-X-C chemokine receptor type 4.

**Supplementary Figure S7**

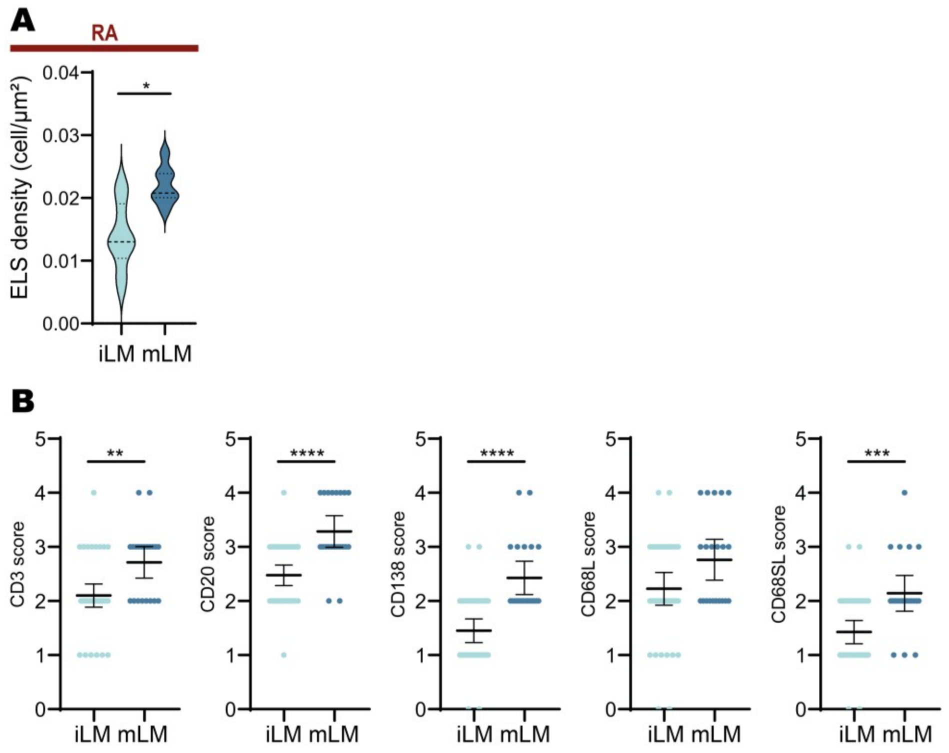


**Fig. S7.** ***Associated with main Figure 4.*** **(A)** Violin plots representing the distribution, mean and quartiles values of ectopic lymphoid structures (ELS) density in cells/µm$^2$ in rheumatoid arthritis (RA) lympho-myeloid (LM) tissues (n= 16 tissues analyzed), assessed using QuPath. p-value was calculated using the Mann-Whitney test, *, p<0.05. **(B)** CD3, CD20, CD138, lining CD68 (CD68L), and sublining CD68 (CD68SL) semi-quantitative scores in lympho-myeloid (LM) synovial tissues from OA patients belonging to the immature LM (iLM, n=40) or mature LM (mLM, n=21) group. p-values were assessed by the Mann-Whithney test, **, p< 0.01; ***, p<0.001; ****, p<0.0001. Individual values, mean and SEM are shown.

**Supplementary Figure S8**

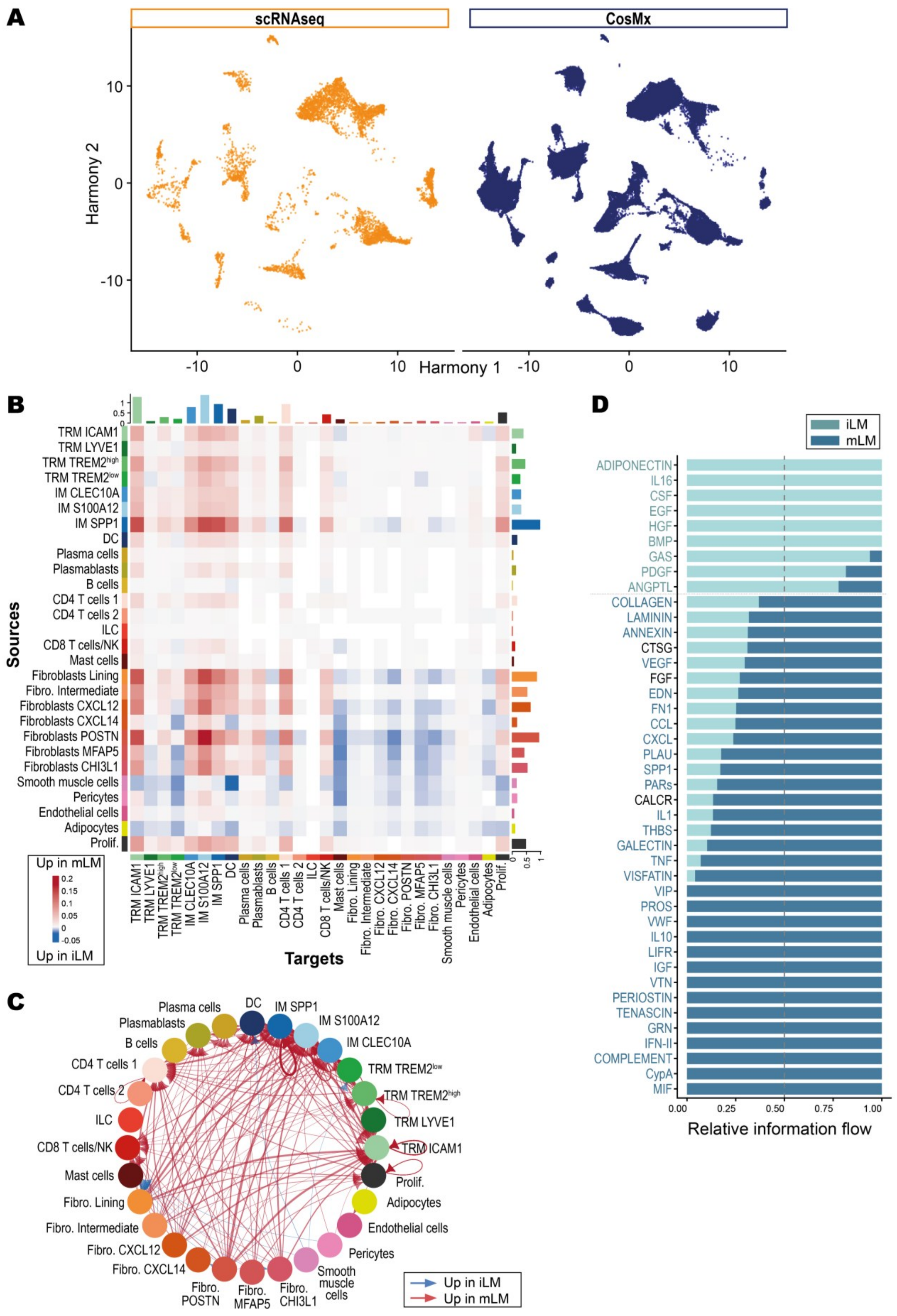

**Fig. S8.** ***Associated with main Figure 5.*** **(A)** Uniform Manifold Approximation and Projection (UMAP) of cells from scRNA-seq (orange) and CosMx (blue) datasets. Each dot represents an individual cell. **(B)** Heatmap illustrating cell-cell communication patterns inferred by CellChat in iLM and mLM synovial tissues. Rows and columns correspond to cell groups (targets and sources, as indicated). Heatmap intensity (red-blue gradient) reflects interaction strength, with red indicating interactions stronger in mLM and blue indicating interactions stronger in iLM. **(C)** Circle plot showing differential cell-cell interaction strength across iLM and mLM synovial tissues. Each node represents a cell population, and the connecting lines indicate interactions. Line color reflects the direction of differential interaction: red for interactions stronger in mLM and blue for interactions stronger in iLM. Line thickness corresponds to the relative interaction strength. **(D)** Bar chart showing the overall information flow of each signaling pathway inferred by CellChat. iLM pathways are shown in light blue and mLM pathways in dark blue. Pathways are ranked based on differences in overall information flow between iLM and mLM. Pathways labeled in black are not significantly different (paired Wilcoxon test). The chart represents the relative contribution of each pathway to intercellular communication in the respective tissue groups.

**Supplementary Figure S9**

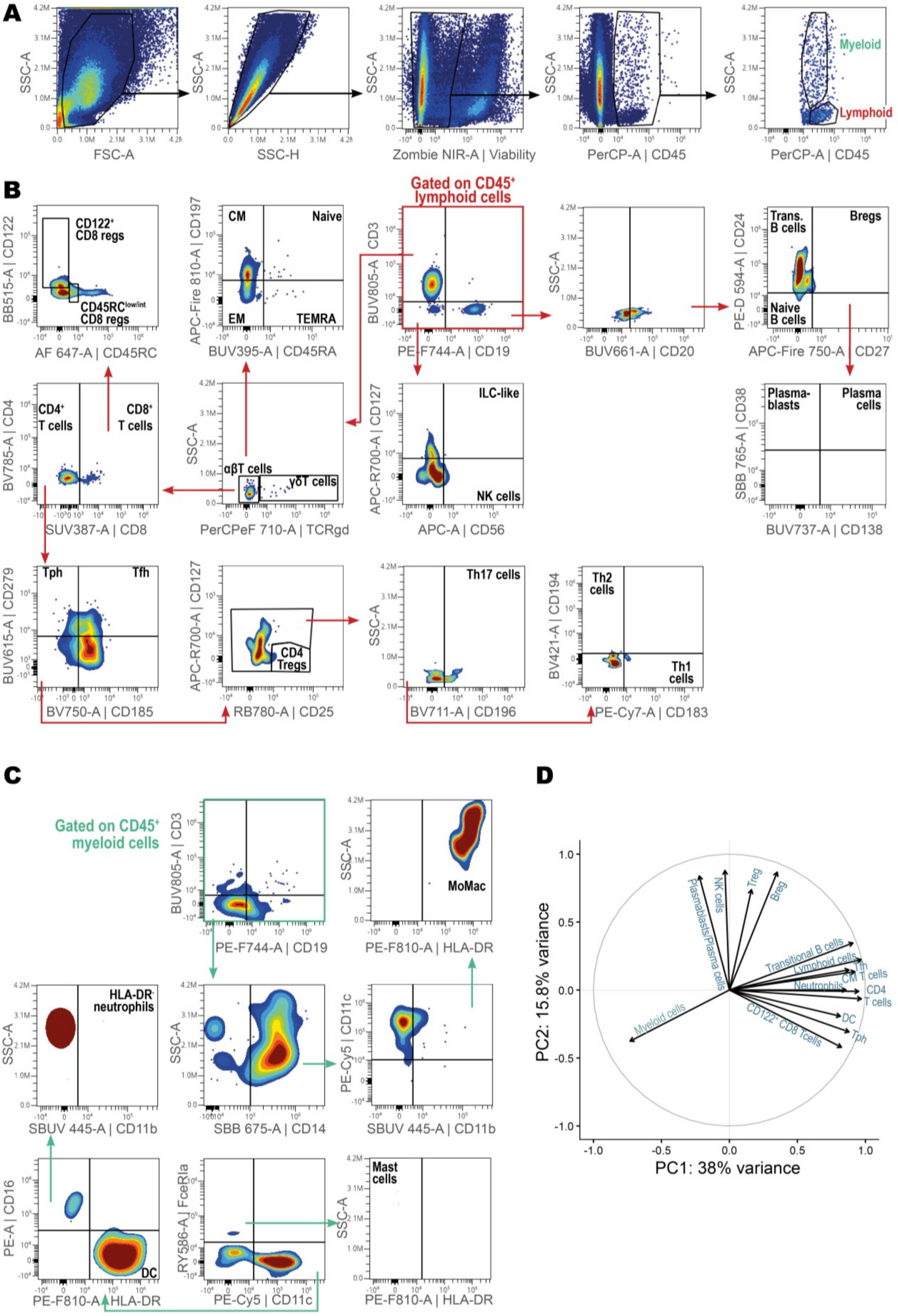

**Fig. S9.** ***Associated with main Figure 5.*** **(A-C)** Gating strategy for spectral cytometry analysis showing forward scatter (FSC) and side scatter (SSC) gating, viability exclusion, and selection of $CD45^{+}$ live cells **(A)**, identification of lymphoid populations, including T cells, B cells, and their subsets, innate lymphoid cells (ILC) and natural killer (NK) cells **(B)**, and identification of myeloid populations, including monocytes/macrophages (MoMac), dendritic cells (DC), and neutrophils **(C)**. **(D)** PCA biplot, the length and direction of the 15 more important variable vectors indicate their influence to PC1 and PC2, which accounts for 38,3% and 15,9% of variance, respectively.

**Supplementary Figure S10**

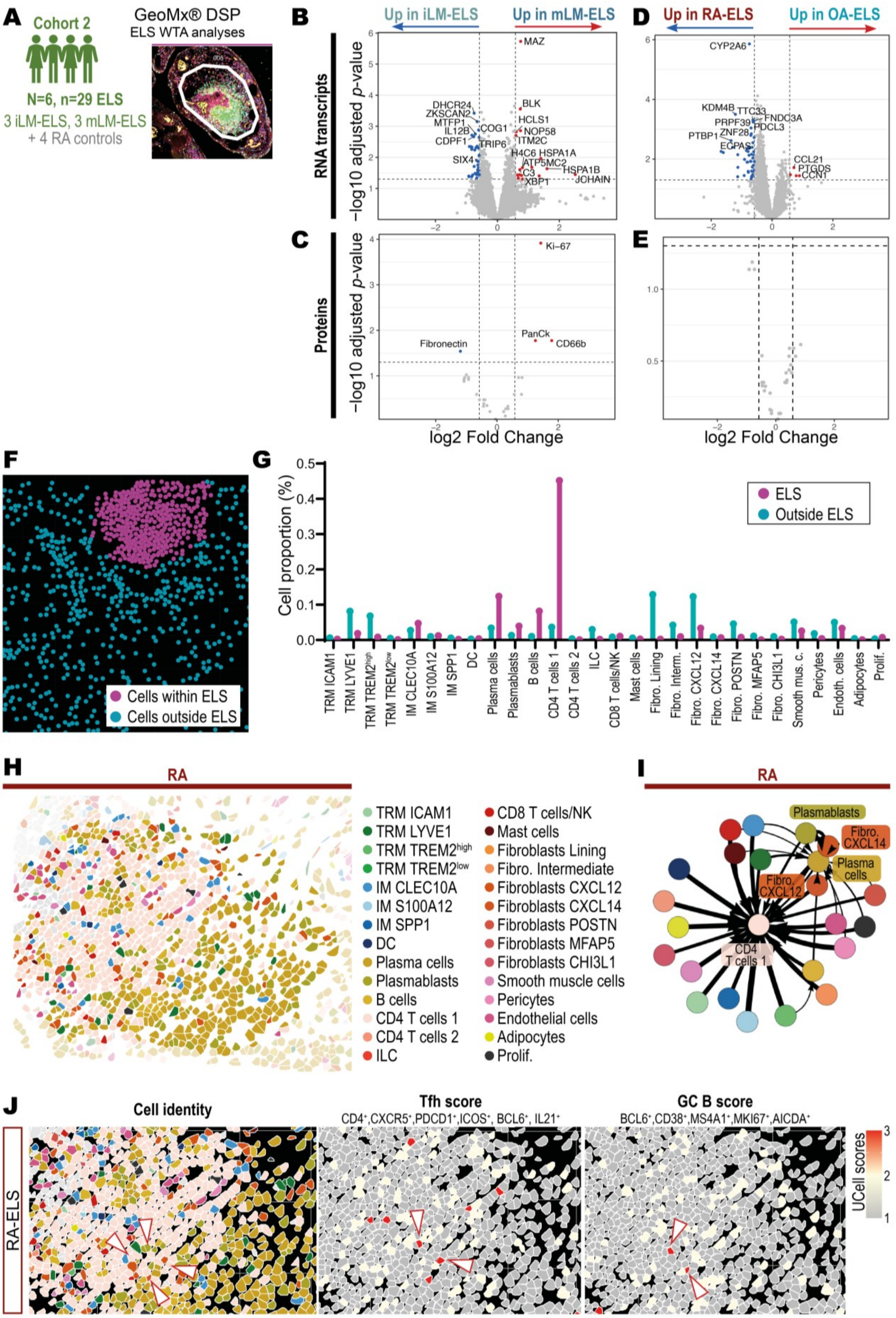

**Fig. S10.** ***Associated with main Figure 6.*** **(A)** For spatial analyses of ELS molecular profile, whole transcriptome atlas (WTA) analyses using the GeoMx Digital Spatial Profiler (DSP) on 6 osteoarthritis (OA) lympho-myeloid (LM) tissues (3 immature LM or iLM and 3 mature LM or mLM, including a total of 29 ectopic lymphoid structures or ELS), and 4 LM rheumatoid arthritis (RA) synovial tissues as controls were used. **(B-E)** Volcano plots presenting differentially expressed genes **(B-D)** and protein **(C-E)** in immature and mature ELS **(B-C)** or in osteoarthritis (OA) and rheumatoid arthritis (RA) ELS **(D-E)** assessed by GeoMx Digital Spatial Imager (DSP). Each dot represents a gene or a protein, the x-axis indicates the –log10(p-adjust) and the y-axis indicates the log2(fold-change) in expression between ROIs. Genes or proteins significantly enriched in immature ELS **(B-C)** or RA **(D-E)** are shown in blue, genes enriched in mature ELS **(B-C)** or OA **(D-E)** are shown in red. **(F)** Representative image illustrating the cell selection strategy using a free-form selection tool for ELS cell attribution ("ELS" cells in fuchsia, "Outside ELS" cells in blue). **(G)** Histogram showing the proportion of each cell type in ELS and outside ELS. **(H)** Representative image of CosMx Image Dimplot representing ELS cell type composition and organization in RA, each cell type is depicted using a different color, according to the corresponding legend. **(I)** Force-directed graph for RA ELS representing directed spatial associations between cell types, derived from the neighborhoods defined by extended Delaunay triangulation (combinatorial radius 4). Each node corresponds to a cell type and is colored accordingly (as specified in the legend in **(H)**). Directed edges (arrows) point from cell type A to cell type B when cells of type B comprise at least 5 % of the neighbors of cells of type A. Edge width reflects the relative weight of these interactions. **(J)** ImageDimPlot showing cells enriched for UCell AddModule scores of follicular helper T cells (Tfh) and germinal center (GC) B cells in RA synovium. Cellular identities are colored according to the legend in **(H)**. Gene signatures used to compute each score are indicated. White arrows highlight cells with pronounced enrichment within ELS.

**Supplementary Table S1**

| | Cohort 1 | | | Cohort 2 | | |
|---|---|---|---|---|---|---|
| | *All* n=186 | *RNA-seq* n=79 | **p-value** (*All* vs. *RNA-seq*) | *All* n=132 | *RNA-seq* n=15 | **p-value** (*All* vs. *RNA-seq*) |
| **Female** % (n) | 68.3% (127) | 74.7% (59) | 0.38; ns | 52.3% (69) | 40.0% (6) | 0.42; ns |
| **Age** years, mean (SD) | 66.5 (10.2) | 66.1 (8.6) | 0.33; ns | 69.7 (8.3) | 68.3 (6.7) | 0.43; ns |
| **Disease duration** years, mean (SD) | ND | ND | ND | 4.4 (4.5) | 5.0 (3.1) | 0.19; ns |
| **BMI**, mean (SD) | ND | ND | ND | 30.3 (5.6) | 29.0 (4.1) | 0.39; ns |

**Table. S1. Characteristics of patients included in Cohort 1 and Cohort 2.** A sub-selection of tissues from patients included in each cohort was analyzed by bulk RNA sequencing (RNA-seq). ND, not determined; ns, not significant; n, number; SD, standard deviation; BMI, body mass index. P-values were calculated using either the Mann–Whitney U test, unpaired Student's t test, or Fisher's exact test, according to sample size and variable type.

**Supplementary Table S2**

| Target | Fluorochrome | Clone | Supplier | Reference | Dilution |
|---|---|---|---|---|---|
| CD8 | Spark UV 387 | SK1 | Biolegend | 344776 | 1/80 |
| CD45RA | BUV395 | HI100 | BD | 568712 | 1/320 |
| CD11b | StarBright UV 445 | 5C6 | Bio-Rad | MCA711SBUV445 | 1/20 |
| CD279 | BUV615 | EH12.1 | BD | 612991 | 1/40 |
| CD20 | BUV661 | 2H7 | BD | 749952 | 1/80 |
| CD138 | BUV737 | MI15 | BD | 612834 | 1/20 |
| CD3 | BUV805 | OKT3 | BD | 750970 | 1/320 |
| CD194 | BV421 | 1G1 | BD | 562579 | 1/20 |
| CD196 | BV711 | G034E3 | Biolegend | 353436 | 1/20 |
| CD185 | BV750 | RF8B2 | BD | 569501 | 1/40 |
| CD4 | BV785 | RPA-T4 | Biolegend | 300553 | 1/20 |
| CD38 | StarBright Blue 765 | AT13/5 | Bio-Rad | MCA1019SBB765 | 1/50 |
| CD122 | BB515 | Mik-?3 | BD | 566059 | 1/20 |
| CD45 | PerCP | 2D1 | Biolegend | 368505 | 1/40 |
| CD14 | StarBright Blue 675 | TuK4 | Bio-Rad | MCA1568SBB675 | 1/20 |
| TCRγδ | PerCP-eFluor 710 | B1.1 | ThermoFisher | 46-9959-41 | 1/50 |
| CD25 | RB780 | 2A3 | BD | 568688 | 1/40 |
| CD16 | PE | 3G8 | Biolegend | 302007 | 1/160 |
| FcεRIα | RealYellow 586 | AER-37 | BD | 753464 | 1/40 |
| CD24 | PE/Dazzle 594 | ML5 | Biolegend | 311134 | 1/40 |
| CD11c | PE-Cy5 | B-ly6 | BD | 561692 | 1/10 |
| CD19 | PE/Fire 744 | HIB19 | Biolegend | 390405 | 1/160 |
| CD183 | PE-Cy7 | G025H7 | Biolegend | 353720 | 1/40 |
| HLA-DR | PE/Fire 810 | L243 | Biolegend | 307683 | 1/160 |
| CD56 | APC | HCD56 | Biolegend | 318310 | 1/20 |
| CD45RC | Alexa Fluor 647 | MT2 | BD | 565857 | 1/160 |
| CD127 | APC-R700 | HIL-7R-M21 | BD | 565185 | 1/20 |
| Viability | Zombie NIR | / | Biolegend | 423105 | 1/4000 |
| CD27 | APC/Fire 750 | O323 | Biolegend | 302846 | 1/20 |
| CD197 | APC/Fire 810 | G043H7 | Biolegend | 353264 | 1/20 |

**Table. S2. List of antibodies used for spectral flow cytometry.** Target antigen, associated fluorochrome, clone, supplier, catalog reference and dilutions are indicated for each antibody.

**Supplementary Table S3**

| Step | Reagent | Duration | Temperature |
|---|---|---|---|
| **Viability staining** | Zombie NIR | 20 minutes | Room temperature (RT) |
| Wash (x3) in PBE buffer (0,1% Bovine Serum Albumin (Sigma Aldrich, A8412) and 2mM EDTA (Sigma Aldrich, E7889) in 1X PBS (Gibco, 20012-019))<br>Centrifugation at 800g, 2 minutes, RT | | | |
| **CD197 staining** | CD197 APC/Fire 810<br>True-Stain Monocyte Blocker (Biolegend,353264, 1/20)<br>Purified NA/LE Human BD Fc block (BD, 564220, 1/40) | 10 minutes | RT |
| **TCRγδ staining** | TCRγδ/PerCP-eFluor 710 | 10 minutes | RT |
| **Chemokine and CD138 staining** | CD138 bUV737<br>CD183 PE-Cy7<br>CD185 BV750<br>CD194 BV421<br>CD196 BV711<br>Brilliant Stain Buffer Plus (BD, 566385, 1/10) | 15 minutes | RT |
| **Low expression marker staining** | CD24 PE/Dazzle 594<br>CD25 RB780<br>CD27 APC/Fire 750<br>CD45RA BUV395<br>CD45RC Alexa Fluor 647<br>CD122 BB515<br>CD127 APC-R700<br>CD279 BUV615<br>FcεRIα RealYellow 586<br>HLA-DR PE/Fire 810 | 20 minutes | RT |
| Wash (x3): PBE buffer, centrifugation at 800g, 2 minutes, RT | | | |
| **Lineage staining** | CD3 BUV805<br>CD4 BV785<br>CD8 Spark UV 387<br>CD11b StarBright UV445<br>CD11c PE-Cy5<br>CD14 StarBright Blue 675<br>CD16 PE<br>CD19 PE/Fire 744<br>CD20 BUV661<br>CD38 StarBright Blue 765<br>CD45 PerCP<br>CD56 APC<br>PBE buffer | 20 minutes | RT |
| Wash (x3): 1X PBS buffer, centrifugation at 800g, 2 minutes, RT | | | |
| **Fixation** | 2% paraformaldehyde (PFA) | 15 minutes | 4°C |
| Wash (x3): PBE buffer, centrifugation at 800g, 2 minutes, RT | | | |
| **Tandem stabilizer** | 1X Tandem stabilizer in PBE buffer | Until acquisition | 4°C |

**Table. S3. Spectral flow cytometry staining procedure.** Each reagent, incubation duration, and temperature are indicated for each step.
